# DRP1 inhibition confers cardioprotection against doxorubicin while preserving anticancer efficacy

**DOI:** 10.64898/2026.02.15.705503

**Authors:** Yali Deng, Sebastian Bass-Stringer, Simon T. Bond, Jonathon Cross, Jia Q Truong, Leonardo M. Hugen, Hei-Yi H. Woo, Ayeshah A. Rosdah, Anne M. Kong, Christopher D. Hart, Kylie L. Gorringe, Rebecca H Ritchie, Elaine Sanij, Brian G. Drew, David W. Greening, Doan Ngo, Jarmon G. Lees, Jessica Holien, Shiang Y. Lim

**Affiliations:** St Vincent’s Institute of Medical Research, Fitzroy, Victoria 3065, Australia; Department of Medicine and Surgery, University of Melbourne, Parkville, Victoria 3010, Australia; Baker Heart and Diabetes Institute, Melbourne, Victoria 3004, Australia; Baker Department of Cardiometabolic Health, University of Melbourne, Melbourne, Victoria 3010, Australia; School of Translational Medicine, Monash University, Melbourne, Victoria 3004, Australia; STEM College, RMIT University, Melbourne, Victoria 3000, Australia; Department of Pharmacology, Faculty of Medicine, Universitas Sriwijaya, Palembang 30126, Indonesia; St Vincent’s Hospital Melbourne, Fitzroy, Victoria 3065, Australia; Peter MacCallum Cancer Centre and the Sir Peter MacCallum Dept of Oncology, University of Melbourne, Melbourne, Victoria 3000, Australia; Drug Discovery Biology, Faculty of Pharmacy and Pharmaceutical Sciences, Monash University, Parkville, Victoria 3052, Australia; Department of Cardiovascular Research, Translation and Implementation, La Trobe University, Melbourne, Victoria 3086, Australia; School of Biomedical Sciences and Pharmacy, University of Newcastle, Callaghan, New South Wales 2308, Australia; Hunter Medical Research Institute, University of Newcastle and Calvary Mater Newcastle, Newcastle, New Lambton Heights, New South Wales 2305, Australia; National Heart Research Institute Singapore, National Heart Centre, 5 Hospital Drive, 169609, Singapore

**Keywords:** Drp1, Mitochondrial fission, Cancer, Doxorubicin-induced cardiotoxicity, Human induced pluripotent stem cells, Cardiac microtissues

## Abstract

**Background:** Anthracyclines such as doxorubicin are effective chemotherapeutics but are limited by cardiotoxicity driven in part by mitochondrial dysfunction. Dysregulated mitochondrial dynamics, particularly excessive dynamin-related protein-1 (Drp1)-mediated fission, contribute to doxorubicin-induced cardiac injury and support selective survival of cancer cells.

**Objectives:** To determine whether DRP1i2, a novel small molecule Drp1 inhibitor targeting a conserved domain shared between human and mouse, can function as a cardio-oncology therapeutic by reducing doxorubicin-induced cardiotoxicity while maintaining or enhancing anti-cancer efficacy.

**Methods:** Cardioprotective effects of DRP1i2 were evaluated in a murine model of chronic doxorubicin cardiotoxicity and in human induced pluripotent stem cell-derived cardiac microtissues exposed to acute doxorubicin injury. Anticancer activity was assessed across multiple cancer cell lines using 2D monolayers and 3D microtissues.

**Results:** *In vivo*, DRP1i2 preserved left ventricular ejection fraction, reduced interstitial fibrosis and cardiomyocyte atrophy, and attenuated doxorubicin-induced myocardial proteomic remodelling. In human cardiac microtissues, DRP1i2 improved viability and restored contractile function despite persistent mitochondrial oxidative stress. DRP1i2 showed modest anticancer activity in MG63 osteosarcoma cells in both 2D and 3D systems and did not diminish doxorubicin efficacy in other cancer models (MDA-MB-231 breast, OVCAR3 ovarian, and A549 lung adenocarcinoma). Combined treatment further enhanced cytotoxicity selectively in MG63 cells.

**Conclusions:** DRP1i2 exerts complementary cardioprotective and anticancer actions through modulation of shared mitochondrial pathways, identifying Drp1 as a druggable target in cardio-oncology. These findings support DRP1i2 as a first-in-class Drp1 inhibitor and highlight mitochondrial dynamics as a promising therapeutic axis to preserve anthracycline efficacy while reducing cardiotoxicity.

**Clinical Perspectives:** Excessive Drp1-mediated mitochondrial fission links anthracycline cardiotoxicity with cancer cell survival. Inhibition with DRP1i2 preserved cardiac structure and function in a chronic doxorubicin cardiotoxicity model without compromising anti-cancer activity, representing mechanism-based cardioprotection, where the heart is protected by directly targeting the molecular processes driving injury. Translation will require pharmacologic profiling and testing in tumour-bearing and comorbid models, followed by early-phase trials to confirm safety and efficacy.

## Introduction

Anthracyclines such as doxorubicin remain indispensable in the treatment of many cancers, yet their therapeutic value is limited by cumulative, dose-related cardiac injury (1). As cancer survival rates improve, a growing number of individuals previously exposed to anthracyclines face an increased risk of heart failure and other cardiovascular complications (2). Current management strategies rely on dose limitation and reactive treatment of heart failure using standard cardiology agents, including angiotensin-converting enzyme inhibitors, angiotensin receptor blockers, beta-blockers, and mineralocorticoid receptor antagonists. Dexrazoxane, a small-molecule iron chelator, is the only cardioprotective agent approved by the United States Food and Drug Administration for reducing doxorubicin-related cardiac injury. This approach is used only in selected high-risk settings because of concerns about potential interference with anticancer efficacy and restrictive labelling. Moreover, dexrazoxane provides only partial protection and is not suitable for all patients (3,4). These limitations underscore an ongoing need for more effective and widely applicable strategies to preserve cardiac health in individuals receiving anthracycline therapy.

Mitochondrial dysfunction is central to doxorubicin-induced cardiotoxicity, converging on impaired bioenergetics, excessive reactive oxygen species (ROS) generation, and activation of cell death pathways (5-7). Mitochondrial dynamics, governed by the balance between fusion and fission, are increasingly recognised as key regulators of these processes (8). Dynamin-related protein 1 (Drp1), a GTPase that drives mitochondrial fission, is activated in response to cellular stress. Excessive Drp1-mediated mitochondrial fission has been implicated not only in cardiomyocyte injury, but also in cancer cell survival, metabolic adaptability, and chemoresistance (5,7). This dual impact makes Drp1 a compelling, yet underexplored, therapeutic target at the intersection of oncology and cardiology, with the potential to address two major clinical challenges simultaneously.

To date, small molecule inhibitors of human Drp1 suitable for translational development have been lacking, and the effects of Drp1 modulation across both cardiac and cancer systems remain insufficiently defined (7,9). The present work addresses this gap by characterising DRP1i2, a novel small molecule inhibitor targeting the human Drp1 GTPase domain. The effects of DRP1i2 are systematically evaluated across complementary models of cancer and cardiac injury, including 2D monolayers and 3D spheroids of human cancer cells, human induced pluripotent stem cell (iPSC)-derived cardiomyocytes and multicellular cardiac microtissues, and *in vivo* murine model of doxorubicin-induced cardiotoxicity. This study aims to define the therapeutic potential of DRP1i2 and to support the development of approaches that enable continued, selective, and effective cancer therapy while better protecting cardiovascular health.

## Methods

### Animal ethics declaration

All experimental procedures were approved by the St Vincent’s Hospital Animal Ethics Committee (AEC No. 003/23) and conducted in accordance with the Australian National Health and Medical Research Council Code for the care and use of laboratory animals, Directive 2010/63/EU, and NIH guidelines. Animal research is reported in accordance with ARRIVE 2.0 guidelines to ensure transparent and reproducible study design, conduct, and reporting.

### *In vivo* model of chronic doxorubicin-induced cardiotoxicity

To induce chronic doxorubicin-induced cardiotoxicity, Male C57BL/6J mice (male, six weeks old) were given weekly intraperitoneal injections of doxorubicin at 5 mg/kg for five consecutive weeks. Briefly, after 7 days of acclimatisation, mice were randomly assigned into four experimental groups (n=6 per group) to receive five-weekly intraperitoneal injections of either vehicle control (0.1% DMSO in saline), DRP1i2 (1 mg/kg), doxorubicin (Dox, 5 mg/kg) or Dox+DRP1i2. Drugs were administered following echocardiographic assessment, and mice were maintained under deep anaesthesia throughout the procedure. Survival and body weight were monitored for 35 days. On day 35, after the final echocardiographic assessment, mice were maintained under deep anaesthesia and humanely euthanized by surgical removal of the heart for subsequent histological and proteomic analyses. Heart weight and tibial length were recorded at the time of sacrifice. All animal experimental procedures were performed by experienced animal surgeons.

### Cardiac microtissue

Cardiomyocyte-only microtissues and multicellular cardiac microtissues were constructed using previously published protocols with modifications (10). Cardiomyocyte-only microtissues (7.5 × 10^4^ cells per microtissue) and multicellular cardiac microtissues (7.5 × 10^4^ total cells per microtissue; 75% cardiomyocytes, 20% endothelial cells, 2.5% cardiac fibroblasts, and 2.5% vascular smooth muscle cells) were treated for 3 days with vehicle control (0.1% DMSO), DRP1i2 (10, 50, or 100 μM), doxorubicin alone, or a combination of doxorubicin and DRP1i2.

### Human cancer cell culture

MDA-MB-231, A549, OVCAR3, and MG63 cells in 2D, and MG63 osteosarcoma 3D spheroids (2 × 10^4^ cells/spheroid) were treated for 5 days with vehicle (0.1% DMSO), DRP1i2 (50 μM), doxorubicin (0.05 μM), or a combination of doxorubicin and DRP1i2.

### Statistical analysis

Data are expressed as mean ± SEM. Statistical analyses were performed using GraphPad Prism. Differences among groups were analysed by unpaired Student’s *t*-test, one-way or two-way ANOVA, as appropriate. When ANOVA indicated a significant effect, *post hoc* multiple-comparison tests were applied as specified in the figure legends. A *P* value < 0.05 was considered statistically significant.

For detailed methods, see the Supplementary Material.

## Results

### DRP1i2 is a small molecule inhibitor of human Drp1

DRP1i2 was identified from our previously reported virtual screen using OpenEye software (11). It is a tryptophan analogue comprising an imidazole-pyridine fused ring at one terminus and an indole ring at the other, with a primary amine attached to its chiral carbon center. DRP1i2 exhibited dose-dependent binding to human Drp1 isoform 3, with an average dissociation constant of 43.03 ± 3.06 μM across six independent experiments (**Figure 1A**). Moreover, it significantly inhibited GPT hydrolysis in a Drp1 GTPase activity assay (**Figure 1B**).

**Figure 1.**
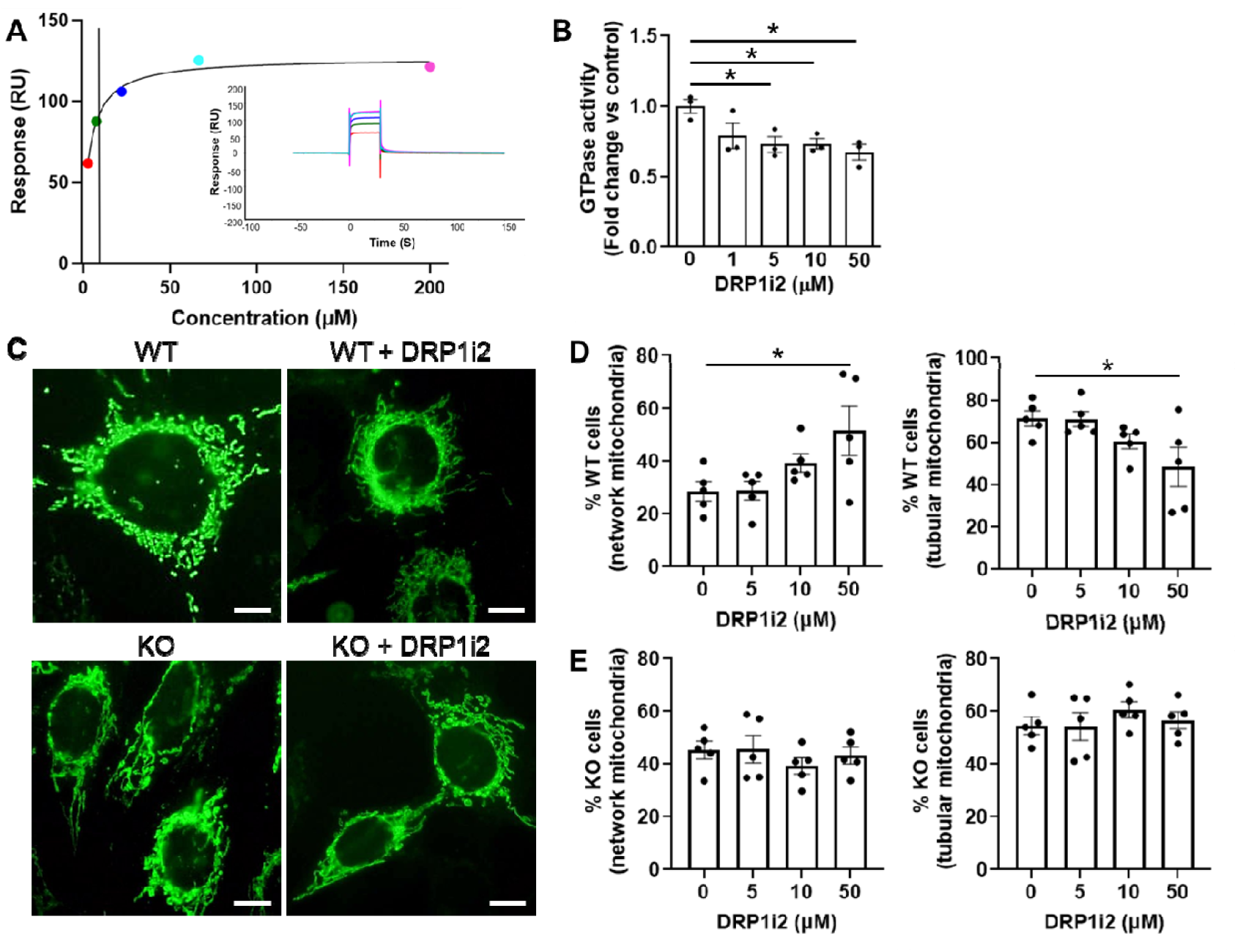
Biochemical and Cellular Characterization of DRP1i2. **(A)** Representative surface plasmon resonance (SPR) sensorgrams showing fitted binding curves and corresponding raw binding responses for DRP1i2 interaction with human Drp1. **(B)** DRP1i2 significantly inhibits the GTPase activity of human Drp1 (n = 3 replicates). **(C)** Representative images of mitochondrial morphology in Drp1 wild-type (WT) and Drp1 knockout (KO) mouse embryonic fibroblasts (MEFs). **(D-E)** Quantification of mitochondrial morphology in Drp1 wild-type **(D)** and Drp1 knockout **(E)** MEFs treated with DRP1i2 (n = 5 replicates). *P<0.05 by one-way ANOVA.

To assess the functional impact of DRP1i2 on Drp1-mediated mitochondrial fission, mitochondrial morphology was analysed in Drp1 wild-type and knockout mouse embryonic fibroblasts (MEFs) treated with DRP1i2. At baseline, Drp1 knockout MEFs exhibited a higher proportion of cells with networked mitochondria compared with wild-type MEFs. DRP1i2 induced a dose-dependent increase in the proportion of wild-type MEFs displaying networked mitochondria, accompanied by a concomitant decrease in cells with tubular mitochondria, reaching statistical significance at 50 μM (**Figure 1C**), consistent with its binding affinity to recombinant Drp1. Importantly, DRP1i2 had no significant effect on mitochondrial morphology in Drp1 knockout MEFs, indicating that its effects are Drp1-dependent (**Figure 1D**). These results suggest that Drp1i2 is a modest but selective Drp1 inhibitor.

### DRP1i2 preserved cardiac function and tissue architecture in a mouse model of chronic doxorubicin-induced cardiotoxicity

C57BL/6J mice received weekly intraperitoneal injections of vehicle, DRP1i2, doxorubicin, or doxorubicin plus DRP1i2 for five weeks (**Figure 2A**). Vehicle- and DRP1i2-treated mice showed progressive weight gain over 35 days, whereas doxorubicin-treated mice, with or without DRP1i2, failed to gain weight and had significantly lower body weight at day 35 compared with vehicle controls (**Figure 2B**). DRP1i2 did not affect doxorubicin-associated body weight changes. Doxorubicin significantly reduced heart weight, tibial length, and the heart weight/tibial length ratio. DRP1i2 alone had no effect on these parameters but prevented doxorubicin-induced reductions in heart weight, tibial length, and normalised heart weight, restoring values to near control levels **(Figure 2C-E)**.

**Figure 2.**
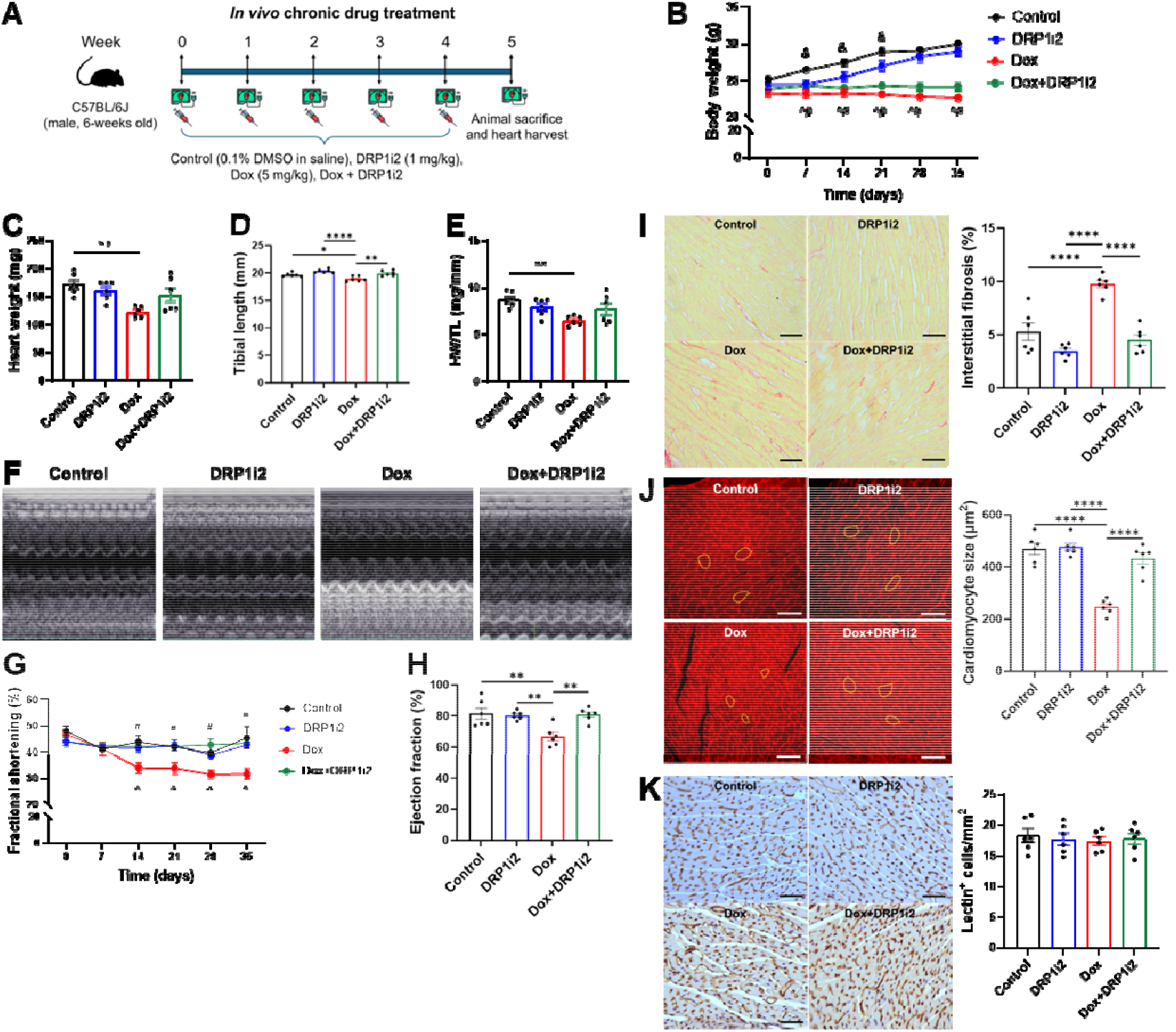
Cardioprotective effect of DRP1i2 in a murine model of chronic doxorubicin-induced cardiotoxicity. **(A)** Treatment regimen in a murine model of doxorubicin (Dox)-induced cardiotoxicity. **(B)** Body weight monitored over 35 days. **(C)** Heart weight, **(D)** tibial length, and **(E)** heart weight/ tibial length (HW/TL) recorded on day 35. **(F)** Representative 2D motion-mode echocardiography images. **(G)** Fractional shortening monitored over 35 days. **(H)** Ejection fraction on day 35. **(I-K)** Representative images and quantification of interstitial fibrosis **(I)**, cross-sectional area of cardiomyocytes **(J)**, and myocardial vascular density **(K)**. Scale bar = 50 μm. n = 6 mice per group (each data point represents an individual mouse). ^&^P<0.05 for DRP1i2 versus Control, ^P<0.05 for Dox versus Control, and ^#^P<0.05 for Dox+DRP1i2 vs. Dox by two-way ANOVA. *P<0.05 and **P<0.01, and ****P<0.0001 by one-way ANOVA

Echocardiography showed progressive doxorubicin-induced cardiac dysfunction, with significant reductions in fractional shortening and ejection fraction from day 14 that continue to decline through day 35. Co-treatment with DRP1i2 preserved cardiac function, preventing reductions in both measures (**Figure 2F-H**). Histologically, DRP1i2 alone did not affect interstitial fibrosis, cardiomyocyte size, or myocardial vascular density. Doxorubicin markedly increased interstitial fibrosis and reduced cardiomyocyte size, both of which were significantly attenuated by DRP1i2 co-treatment. Myocardial vascular density was unchanged across all groups (**Figure 2I-K**).

### DRP1i2 mitigated doxorubicin-induced alterations in cardiac proteome homeostasis

Mass spectrometry-based proteomics of mouse heart tissues identified a total of 4391, 4287, 4264 and 4264 proteins in the vehicle control, DRP1i2, doxorubicin, and Dox+DRP1i2 groups, respectively, with high reproducibility (CV <25%) and consistent intensity distributions (**Supplementary Fig. 1A-C, Supplementary Tables 1-2**). Principal component analysis demonstrated a clear doxorubicin-induced proteomic shift, partial overlap of DRP1i2 with the doxorubicin cluster, and a distinct Dox+DRP1i2 profile (**Figure 3A**). A total of 3,788 proteins were commonly identified across all groups, with smaller subsets uniquely detected in each condition (**Figure 3B**).

**Figure 3.**
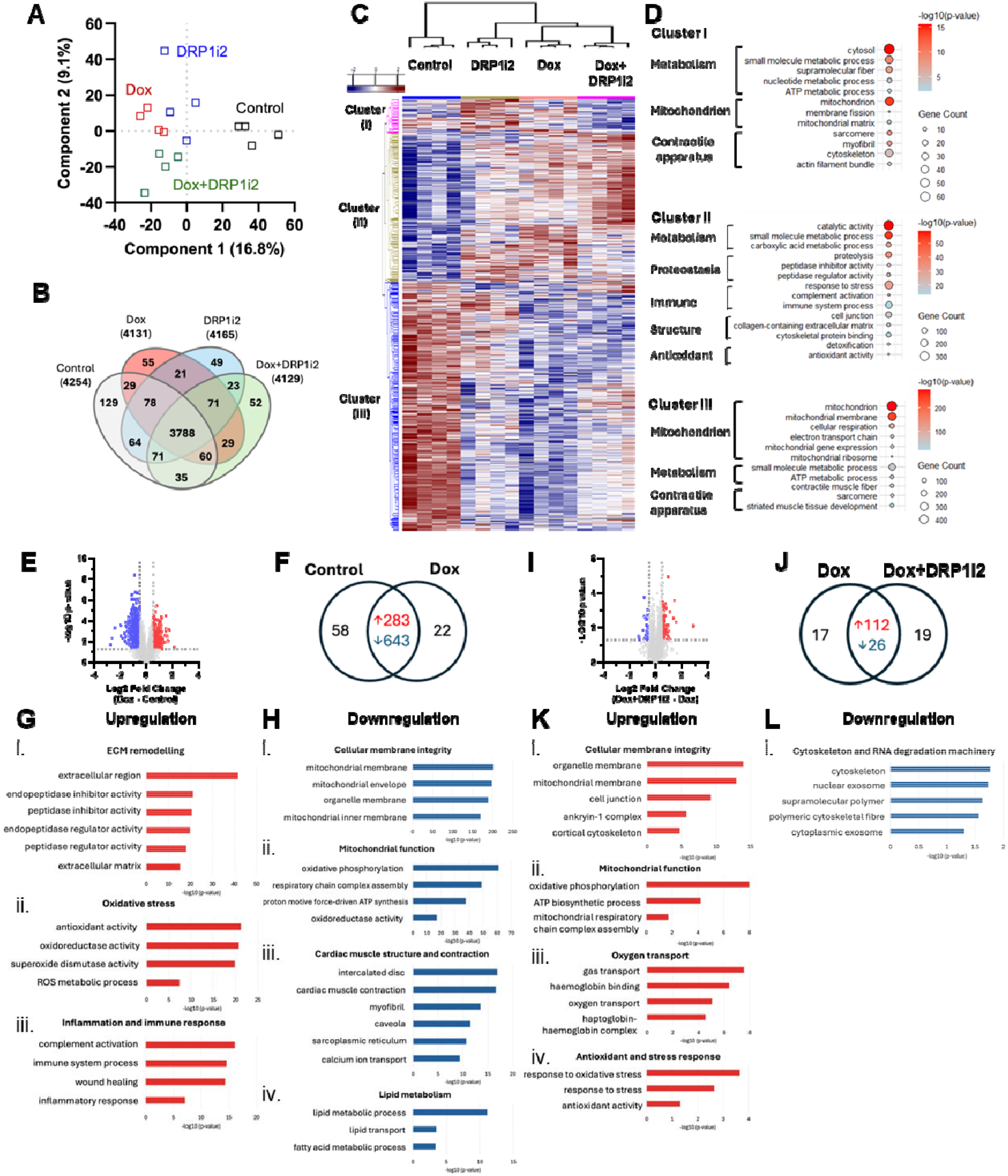
Global proteome profiling of murine heart tissue in response to doxorubicin-induced cardiotoxicity and DRP1i2 treatment. (**A**) Principal component analysis of mouse heart tissues. **(B)** Venn diagram of the number of proteins identified in each group (n = 4 moue heart samples per group, with stringent protein identification and protein/peptide level FDR of <0.01). (**C**) Clustered heatmap of 1999 significant proteins identified by one-way ANOVA analysis (P<0.05), with z-scores normalised and a minimum cutoff of 3 out 4 replicates. **(D)** Balloon plots showing significantly enriched Gene Ontology terms for selected clusters, identified using g:Profiler. Pathways are ranked by –log10 p-value, with cluster-specific significance thresholds applied (cluster i: >2; cluster ii: >10; cluster iii: >20). (**E–H)** Differential protein expression and Gene Ontology (GO) enrichment in mouse hearts treated with doxorubicin (Dox) versus vehicle. **(E)** Volcano plot showing significantly downregulated (blue; p < 0.05, log□FC < –0.5) and upregulated proteins (red; p < 0.05, log□FC > 0.5). **(F)** Venn diagram showing significantly regulated and unique proteins. GO enrichment (g:Profiler) for proteins upregulated and unique to Dox **(G)** and for proteins downregulated in Dox and unique to vehicle **(H). (I–L)** Differential protein expression and GO enrichment in Dox+OB37 versus Dox. **(I)** Volcano plot showing significantly downregulated (blue) and upregulated proteins (red) using the same statistical thresholds. **(J)** Venn diagram showing significantly regulated and unique proteins. GO enrichment for proteins upregulated and unique to Dox+OB37 **(K)** and for proteins downregulated in Dox+OB37 and unique to Dox **(L)**.

Using stringent inclusion criteria (protein identification within each group and FDR-controlled protein/peptide stringency), differential expression analysis with Gene Ontology enrichment identified three major clusters reflecting treatment-specific biological responses (**Figure 3C, Supplementary Table 3**). *Clusters I* and *III* were strongly enriched for mitochondrial- and cardiac-associated proteins, indicating a central role for mitochondrial regulation in the response. *Cluster I*, upregulated by DRP1i2 alone and reduced in Dox+DRP1i2, was significantly enriched for myofibrillar and cytoskeletal organization, mitochondrial matrix and metabolic pathways, and mitochondrial dynamics (e.g. Drp1) (**Figure 3D, Supplementary Table 3A**). *Cluster II*, induced by doxorubicin, was dominated by oxidative stress, inflammatory, and proteostasis networks, with enrichment of antioxidant activities, immune response and extracellular matrix remodelling, consistent with a robust stress-adaptive response to doxorubicin-induced cardiac injury (**Figure 3D, Supplementary Table 3B**). *Cluster III*, suppressed by doxorubicin and partially restored by DRP1i2 co-treatment, was enriched for oxidative phosphorylation, electron transport chain components, mitochondrial dynamics and sarcomere organisation, highlighting preservation of cardiac structural and metabolic integrity with DRP1i2 (**Figure 3D, Supplementary Table 3C**).

Pairwise comparisons further defined treatment-specific protein network alternations. DRP1i2 alone produced a mild adaptive remodelling signature (**Supplementary Figure 2A-D, Supplementary Table 4A-D**). Doxorubicin induced extensive remodelling, with 283 proteins significantly upregulated and 414 downregulated in expression relative to vehicle control (**Figure 3E-F, Supplementary Table 5A-B**). Upregulated pathways included extracellular matrix remodelling, protease regulation, oxidative stress responses, and immune activation (**Figure 3Gi-iii, Supplementary Table 5C**). Downregulated proteins were enriched for mitochondrial inner membrane components, respiratory chain complexes, ATP synthesis pathways, excitation-contraction coupling proteins, and fatty acid β-oxidation enzymes (**Figure 3Hi-iv, Supplementary Table 5D**), indicating that doxorubicin caused pronounced downregulation of proteins critical for mitochondrial integrity, energy metabolism, and cardiac contractile function.

DRP1i2 co-treatment partially restored the doxorubicin-altered proteome, increasing 112 proteins and reducing 26 proteins, including a distinct subset uniquely associated with cardioprotection (**Figure 3I-J, Supplementary Table 6A-B**). Restored pathways included membrane stability and cytoskeletal anchoring (**Figure 3Ki, Supplementary Table 6C**), oxidative phosphorylation and respiratory complex assembly (**Figure 3Kii, Supplementary Table 6C**), and oxygen transport (e.g. HBA, HBB, HP) (**Figure 3Kiii, Supplementary Table 6C**). In addition, antioxidant and stress-response proteins, including MGST1, TXNIP, GCLM, and ROMO1, were upregulated (**Figure 3Kiv, Supplementary Table 6C**). Proteins downregulated by DRP1i2 co-treatment were primarily associated with cytoskeletal remodelling, such as BYSL, NEFM, PDE4DIP, MMS19, and NFEL, as well as RNA degradation proteins (**Figure 3Li, Supplementary Table 6D**).

Together, these findings in cardiac tissue demonstrated that DRP1i2 mitigates doxorubicin-induced disruptions in mitochondrial function, cytoskeletal integrity, and oxidative stress pathways, supporting its role as a cardioprotective modulator of cardiac proteome homeostasis.

### DRP1i2 reduced doxorubicin-induced cardiotoxicity in human cardiac microtissues

To translate *in vivo* findings to a human context, the effects of DRP1i2 were examined in engineered human cardiac microtissues. Immunocytochemistry confirmed that cardiomyocyte[only microtissues consisted exclusively of cTnT□ cardiomyocytes (**Figure 4A**), whereas multicellular microtissues contained the expected heterogeneous population of CD31□ endothelial cells, vimentin□ fibroblasts, and SM22□α□ smooth muscle cells organised around cardiomyocytes (**Figure 4B**).

**Figure 4.**
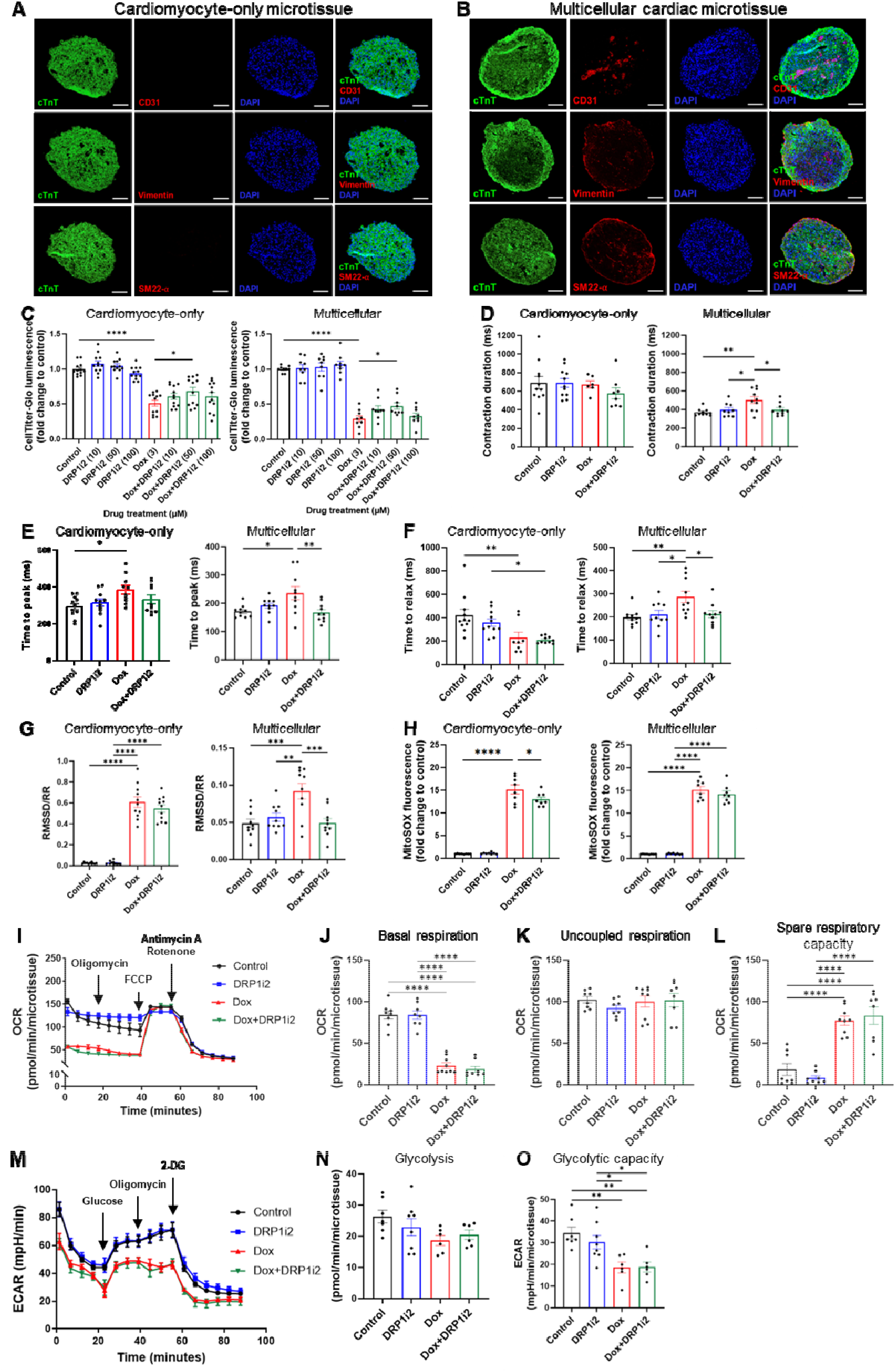
DRP1i2 protects human iPSC-derived cardiac microtissues from doxorubicin-induced cytotoxicity. **(A-B)** Immunocytochemistry of cardiomyocyte-only **(A)** and multicellular **(B)** cardiac microtissues stained for cTnT (cardiomyocytes), CD31 (endothelial cells), vimentin (fibroblasts), and SM22-α (smooth muscle cells); nuclei counterstained with DAPI. Scale bar = 100□μm. **(C)** Cell viability measured by CellTiter-Glo (n = 9-12 microtissues). **(D–G)** Contractile function, including total contraction duration **(D**), time to peak contraction (E), time to relaxation **(F)**, and beat-to-beat variability (RMSSD/RR) **(G)** (n = 6-11 microtissues). **(H)** Mitochondrial ROS levels assessed by MitoSOX fluorescence (n = 8 microtissues). **(I–L)** Seahorse analysis of mitochondrial respiration (OCR) in multicellular cardiac microtissues: representative traces **(I)**, basal respiration **(J)**, uncoupled respiration **(K)**, and spare respiratory capacity **(L)** (n = 8-9 microtissues). **(M–O)** Seahorse analysis of glycolytic function (ECAR) in multicellular cardiac microtissues: representative traces **(M)**, glycolysis **(N)**, and glycolytic capacity **(O)** (n = 6-8 microtissues). *P<0.05, **P<0.01, ***P<0.001, ****P<0.0001 by one-way ANOVA.

DRP1i2 (10-100 μM) had no effect on basal viability in either cardiomyocyte-only or multicellular cardiac microtissues, confirming good tolerability under unstressed conditions (**Figure 4C**). Doxorubicin induced a pronounced loss of viability across both models, establishing a robust *in vitro* cardiotoxicity phenotype. DRP1i2 co-treatment significantly attenuated this toxicity, with maximal rescue at 50 μM in both microtissue types (**Figure 4C**). Accordingly, 50 μM was selected for subsequent functional and mechanistic studies. DRP1i2 alone did not alter contractile function (**Figure 4D-G**). In cardiomyocyte-only microtissues, doxorubicin induced prolonged time to peak (**Figure 4E**), accelerated relaxation (**Figure 4F**), and increased beat-to-beat variability (**Figure 4G**), none of which were rescued by DRP1i2. In multicellular microtissues, doxorubicin slowed contraction and relaxation kinetics and increased variability, and DRP1i2 significantly attenuated these abnormalities, restoring function toward control levels (**Figure 4D-G**). This selective rescue indicates that DRP1i2-mediated protection requires multicellular interactions.

Doxorubicin induced a ∼15-fold increase in mitochondrial ROS in both microtissue types (**Figure 4H**). DRP1i2 alone did not alter baseline ROS levels and only modestly attenuated doxorubicin-induced ROS, indicating that its survival benefit occurs despite persistent oxidative stress and likely involves mechanisms that do not fully prevent doxorubicin-mediated mitochondrial injury. (**Figure 4H**). To assess cell-type-specific vulnerability, 2D monocultures were examined (**Supplementary Figure 3**). Doxorubicin impaired viability and mitochondrial function across all cell types, whereas DRP1i2 did not improve viability, reduce mitochondrial ROS, or restore mitochondrial membrane potential (**Supplementary Figure 3**), indicating that its protective effects emerge only in multicellular tissue contexts.

Oxygen consumption and extracellular proton flux analysis of multicellular cardiac microtissues using the Seahorse assay showed that doxorubicin significantly reduced both basal and ATP-linked respiration, as indicated by the oxygen consumption rate (OCR) (**Figure 4I-J, Supplementary Figure 4A**), while uncoupled OCR was preserved (**Figure 4K**), increasing spare respiratory capacity (**Figure 4L**) due to reduced basal respiration. Non-mitochondrial OCR and proton leak were modestly reduced (**Supplementary Figure 3B-C**). Doxorubicin also impaired glycolytic capacity (**Figure 4M, O**) and glycolytic reserve (**Supplementary Figure 3D**), with non-significant reductions in basal glycolysis (**Figure 4N**) and non-glycolytic acidification (**Supplementary Figure 3E**). DRP1i2 did not alter OCR or ECAR when administered alone, nor did it affect the changes induced by doxorubicin.

Together, these findings show that doxorubicin suppresses mitochondrial oxidative metabolism and glycolytic responsiveness in human cardiac microtissues. DRP1i2 improves viability and restores contractile function selectively in multicellular microtissues, despite limited effects on mitochondrial ROS and bioenergetics, highlighting a multicellular mechanism of cardioprotection.

### DRP1i2 exhibits modest anti-cancer activity without compromising doxorubicin efficacy in MG63 osteosarcoma cells

Drp1 expression and phosphorylation were profiled across four cancer cell lines using lineage-matched non-tumorigenic controls (**Figure 5A-C, Supplementary Figure 5**). Total Drp1 was significantly elevated only in MG63 osteosarcoma cells relative to BJ1 fibroblasts (**Figure 5A**). Activating phosphorylation at serine 616 was increased across all cancer lines (**Figure 5B, Supplementary Figure 5**), whereas serine 637 phosphorylation remained low and unchanged (**Figure 5C, Supplementary Figure 5**), indicating selective enhancement of Drp1 activation in malignant cells.

**Figure 5.**
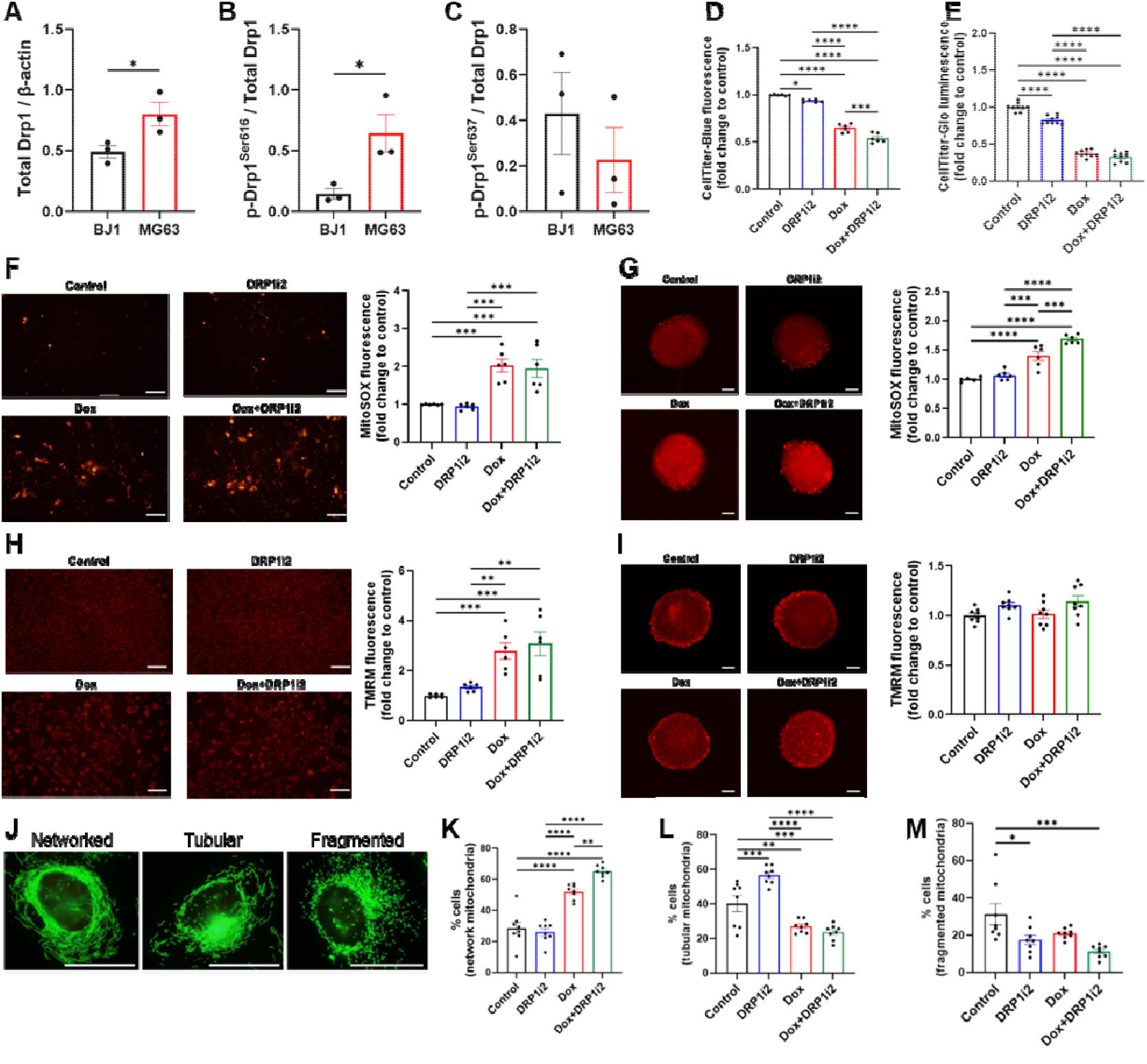
Doxorubicin and DRP1i2 promote MG63 osteosarcoma cancer cell death and mitochondrial dysfunction. **(A-C)** Semi-quantitative analysis of total Drp1 **(A)**, p-Drp1 616 **(B)**, and p-Drp1 637 **(C)** in BJ1 fibroblasts (control) and MG63 osteosarcoma cells (n = 3). **(D-E)** Cell viability of MG63 osteosarcoma cells cultured as 2D monolayers **(D**, n = 6) or 3D spheroids **(E**, n = 9**). (F-G)** Representative images and quantitative analysis of mitochondrial ROS levels in 2D monolayers **(F**, n = 6, scale bar = 100 μm) or 3D spheroids **(G**, n = 6, scale bar = 50 μm**). (H-I)** Representative images and quantitative analysis of mitochondrial membrane potential in 2D monolayers **(H**, n = 6, scale bar = 100 μm) or 3D spheroids **(I**, n = 8 spheroids, scale bar = 50 μm**). (J-M)** Representative images **(J)** and quantification of mitochondrial morphology classified as networked **(K)**, tubular **(L)**, or fragmented **(M)** (n = 8, scale bar = 100 μm). *P < 0.05, **P < 0.01, ***P < 0.001, ****P < 0.0001 by one-way ANOVA.

Despite increased Drp1 serine 616 phosphorylation levels, pharmacological inhibition of Drp1 with DRP1i2 did not reduce viability in MDA-MB-231 (breast cancer), OVCAR3 (ovarian cancer), or A549 (lung cancer) cells (**Supplementary Figure 6**), but modestly decreased viability in MG63 cells in both 2D and 3D culture (**Figure 5D-E**). Doxorubicin reduced viability in all cancer lines, and co-treatment with DRP1i2 further decreased viability only in MG63 cells (**Figure 5D-E, Supplementary Figure 6**). Importantly, DRP1i2 did not diminish doxorubicin efficacy in any model, supporting its compatibility with chemotherapy.

Mechanistically, doxorubicin, but not DRP1i2, significantly increased mitochondrial ROS in MG63 cells (**Figure 5F-G**). Combined treatment further elevated ROS in 3D spheroids but not in 2D cultures (**Figure 5F-G**). Doxorubicin increased mitochondrial membrane potential in 2D cultures only; DRP1i2 alone or in combination had no effect (**Figure 5H-I**). Seahorse analysis of 3D MG63 spheroids revealed no significant changes in mitochondrial respiration or glycolytic activity following treatment with doxorubicin, DRP1i2, or their combination (**Supplementary Figure 7**). Mitochondrial morphology analysis showed predominantly tubular and fragmented mitochondria in vehicle-treated MG63 cells (**Figure 5J-M**). DRP1i2 reduced fragmentation and increased tubular structures, consistent with inhibition of Drp1-mediated fission. Doxorubicin promoted a fused, networked phenotype, and co-treatment with DRP1i2 further enhanced mitochondrial fusion beyond doxorubicin alone (**Figure 5J-M**).

## Discussion

Anthracyclines remain foundational in cancer therapy, yet their clinical utility is constrained by cumulative, dose-dependent cardiotoxicity that manifests as early systolic dysfunction, adverse remodelling, and progressive cardiac atrophy. Despite decades of investigation, effective cardioprotective strategies that preserve oncologic efficacy remain limited. In this study, we identify excessive Drp1-mediated mitochondrial fission as a central, druggable mechanism linking doxorubicin-induced cardiac injury to cancer cell vulnerability, and we demonstrated that DRP1i2, a novel small-molecule inhibitor of Drp1, exhibits therapeutic efficacy as a context-dependent cardio-oncology application. Across complementary *in vivo* and human-relevant *in vitro* systems, DRP1i2 altered the trajectory of anthracycline cardiotoxicity. In a chronic murine model, DRP1i2 preserved left ventricular ejection fraction, reduced interstitial fibrosis, and prevented cardiomyocyte atrophy, hallmarks of early and clinically relevant cardiotoxicity. These functional benefits were mirrored in human iPSC-derived multicellular cardiac microtissues, where DRP1i2 restored contractile performance and improved viability despite persistent mitochondrial oxidative stress. Importantly, DRP1i2 did not attenuate doxorubicin’s anticancer efficacy in various adenocarcinoma cancer models tested and selectively enhanced cytotoxicity in MG63 osteosarcoma cells. Together, these findings position Drp1 inhibition as a viable strategy capable of decoupling cardioprotection from oncologic compromise.

A key translational insight from this study is that DRP1i2-mediated cardioprotection was evident only in physiologically relevant multicellular 3D cardiac microtissues, not in reductionist 2D monocultures. While DRP1i2 failed to rescue isolated cardiomyocytes, it restored contractile function and viability in multicellular constructs containing cardiomyocytes, endothelial cells, fibroblasts, and smooth muscle cells. These findings parallel clinical experience with dexrazoxane and underscores the critical role of tissue architecture, paracrine signalling, and cell-cell interactions in shaping cardiotoxic responses (12,13). Endothelial-derived nitric oxide signalling, extracellular matrix stabilisation, and electromechanical coupling likely contribute to this emergent protection, explaining why contractile recovery occurred despite persistent mitochondrial ROS and why nonspecific antioxidant strategies have failed clinically (14).

### Drp1-mediated fission as an organising mechanism in anthracycline cardiotoxicity

Proteomic and phenotypic analyses indicate that early doxorubicin cardiotoxicity is driven predominantly by atrophic remodelling rather than cardiomyocyte loss (15,16). Coordinated downregulation of sarcomeric and cytoskeletal proteins, activation of proteolytic pathways, suppression of fatty acid oxidation, and impairment of mitochondrial respiratory complexes collectively point to energetic insufficiency as a primary driver of contractile dysfunction (17,18). Excessive Drp1-dependent mitochondrial fission provides a unifying upstream mechanism for these changes, as fragmented mitochondrial networks disrupt cristae architecture, limit ATP production, and amplify localised oxidative stress (7,19). Cardiac microtissue studies further show that Drp1-mediated pathology is not cell autonomous. While doxorubicin markedly increased cell death and mitochondrial oxidative stress, pharmacological Drp1 inhibition with DRP1i2 improved survival without affecting baseline viability. Restoration of contractile function, however, occurred only in multicellular microtissues but not in cardiomyocyte-only constructs, highlighting the importance of multicellular interactions.

Pharmacological inhibition of Drp1 with DRP1i2 disrupted this pathological cascade. DRP1i2 preserved mitochondrial network integrity, stabilised sarcomeric architecture, attenuated fibrotic remodelling, and maintained contractile function despite persistent mitochondrial injury. Doxorubicin-induced ROS and suppression of oxidative and glycolytic flux were only modestly or not rescued, indicating that preservation of mitochondrial network integrity and spatial energy distribution, rather than maximal ATP output, is sufficient to sustain myocardial structure and function under anthracycline stress (20). These data support a revised model of anthracycline cardiotoxicity in which pathological mitochondrial fission acts as an upstream organiser of energetic failure, proteostatic imbalance, oxidative stress, and adverse remodelling.

### Systemic implications and relevance to paediatric cardio-oncology

Beyond the heart, doxorubicin induced a systemic cachectic phenotype, including reductions in body mass and cardiac mass, as well as impaired longitudinal bone growth in young mice (15,16). DRP1i2 prevented tibial shortening, suggesting a potential role for mitochondrial dynamics in systemic growth regulation affected by anthracyclines. While the precise mechanisms remain to be determined, it is possible that preservation of mitochondrial function or redox balance may contribute to maintaining growth plate activity or endocrine signalling during critical developmental windows (21,22). This finding may have particular relevance in paediatric and adolescent cardio-oncology, where long-term growth impairment and cardiotoxicity intersect to influence survivorship outcomes.

### Context-dependent exploitation of Drp1 vulnerability in cancer

Mitochondrial dynamics also regulate cancer cell survival, metabolic plasticity, and therapy resistance (7,23,24). Although elevated Drp1 expression and phosphorylation were observed across multiple cancer cell types, only MG63 osteosarcoma cells exhibited sensitivity to DRP1i2, highlighting that Drp1 dependency is context-specific rather than universal. Because osteosarcoma predominately affects teenagers and young adults, identifying metabolic vulnerabilities such as Drp1 dependence may have particular relevance for improving therapeutic strategies in this younger population (25). MG63 cells displayed high basal Drp1 activity and fragmented mitochondrial networks, consistent with reliance on fission-mediated quality control and metabolic adaptability (26,27).

Compared to doxorubicin alone, DRP1i2 combined with doxorubicin further reduced viability of MG63 cells in 2D cultures but not in 3D spheroids, while it increased mitochondrial ROS in 3D but not in 2D. DRP1i2 alone did not induce oxidative stress or collapse mitochondrial membrane potential. Instead, it promoted mitochondrial elongation, which could potentially influence mitochondrial distribution and turnover. Despite these structural changes, Seahorse analysis of 3D spheroids revealed no significant alterations in mitochondrial respiration or glycolysis with doxorubicin, DRP1i2, or their combination, indicating that basal metabolic flux is largely maintained. When combined with doxorubicin, this structural constraint may contribute to altered mitochondrial responses, with effects that differ between 2D and 3D culture conditions, which model tumour hypoxia, nutrient gradients, and metabolic heterogeneity (28,29). These findings suggest that Drp1 inhibition sensitises select tumours by limiting adaptive mitochondrial remodelling rather than by directly exacerbating oxidative injury.

### Implications for precision cardio-oncology

Collectively, this work establishes excessive Drp1-mediated mitochondrial fission as a shared, targetable mechanism underlying anthracycline cardiotoxicity and cancer cell vulnerability. RP1i2 functioned as a context-dependent modulator, stabilising mitochondrial networks and preserving structure-function coupling in cardiomyocytes, while restricting mitochondrial structural adaptation in cancer cells. The absence of detectable systemic toxicity with DRP1i2 administration further supports the feasibility of Drp1 inhibition as a therapeutic strategy.

These findings advance a precision cardio-oncology paradigm in which mitochondrial dynamics are selectively modulated to protect the heart without compromising, and in select contexts, enhancing anticancer efficacy. They also underscore the necessity of human-relevant multicellular models to uncover clinically concordant cardioprotective mechanisms. Targeting pathological mitochondrial fission emerges as a therapeutic axis for mitigating anthracycline cardiotoxicity while preserving the oncologic intent of treatment.

## Study limitations

This study assessed DRP1i2 in a limited set of cancer models, with the strongest anti-tumour effects in MG63 osteosarcoma cells, which restricts generalisability to other tumour types, molecular subgroups, and chemotherapy regimens. Although 3D cardiac microtissues better recapitulate myocardial architecture than 2D cultures, they lack perfusion, immune cells, and neurohormonal regulation. The *In vivo* studies used a chronic doxorubicin model in healthy, non-tumour-bearing mice, isolating cardiac effects but not capturing tumour-host interactions, concomitant therapies, cardiovascular comorbidities, or other anthracycline regimens used clinically. Finally, DRP1i2 functions as a relatively modest Drp1 inhibitor, which constrains the extent to which full Drp1 suppression could be achieved across experimental settings.

## Supporting information

Supplementary materials (Proteomics)

## Acknowledgements

The authors gratefully acknowledge Dane Newman for providing the A549 lung cancer cell line, which was essential for the experiments conducted in this study.

## Author contributions

YD and SYL conceived the project, designed, and performed the experiments, and analysed and interpreted data. SB, STB, JC, JQT, LMH, HHW, AAR, AMK, and DWG performed experiments and analysed data. AAR, AMK, CH, KLG, DN, RHR, ES, BGD, DWG, JGL, JH, and SYL provided materials and methodologic input. YD, DWG, DN, JGL, JH, and SYL provided interpretative review of study findings. YD drafted the original manuscript and revised the final version. All authors have reviewed, revised, and approved the final version of the manuscript.

## Availability of data

Mass spectrometry-based proteomics data were deposited into the ProteomeXchange Consortium via the MassIVE partner repository and available via MassIVE under the identifier MSV000099244.

