## Supplementary materials for "DRP1 inhibition confers cardioprotection against doxorubicin while preserving anticancer efficacy"

**Supplementary methods**

**Protein expression and purification**

The recombinant human Drp1 protein (His-tagged human Drp1 isoform 3, kindly provided by Professor Michael Ryan, Monash University, Australia) was generated as described previously (1). Briefly, cDNA for human Drp1 (isoform 3) was cloned into pET21b vector and transformed in Rosetta (DE3) competent cells (Novagen, MA, Merck Millipore). Transformed cells were propagated in Luria-Bertani broth and protein expression was initiated by the addition of 0.5 mM isopropyl β-D-1-thiogalactopyranoside. Cells were harvested by centrifugation at 3,500 g for 20 minutes at 4oC, and the pellet was re-suspended in lysis buffer containing 50 mM Tris HCl (pH 7.3), 500 mM NaCl, 50 mM imidazole, 2.5 mM β-mercaptoethanol, 0.01 mM LEUPEP, 0.1 mM AEBSF and 1 mM benzaminidium chloride. The cells were lysed using a pre-cooled EmulsiFlex-C5 homogenizer (Avestin, Ontario, Canada) on ice and clarified by centrifugation at 17,000 g for 30 minutes. The clarified cell lysate was loaded onto a 5 mL Nickel Chelating Sepharose Fast Flow column (GE Healthcare, Buckinghamshire, UK) and Drp1 protein was eluted with wash buffer containing 50 mM Tris-HCl (pH 7.6), 150 mM NaCl, 2.5 mM β-mercaptoethanol and 400 mM imidazole. Eluted proteins were equilibrated with a buffer containing 50 mM Tris-HCl (pH 7.6), 150 mM NaCl, 10% glycerol and 2 mM TCEP using a PD-10 desalting column (GE Healthcare).

**Surface plasmon resonance (SPR)**

All SPR experiments were performed on a Biacore T200 instrument (GE Healthcare Life Sciences, Uppsala, Sweden) at 25°C in the presence of HEPES buffered saline containing 20 mM phosphate (pH 7.5), 137 mM NaCl, 2.7 mM KCl and 0.05% Tween-20 (HBS+P). Protein was tethered to a NiHC500M sensor chip (XanTec bioanalytics, Germany) until approximately 5000RU was achieved. DRP1i2 was purchased from Chembridge (ID 15462333) and dissolved in 100% dimethyl sulfoxide (DMSO) and then added to HBS+P at a final DMSO concentration of 2%. Prior to conducting compound SPR, a DMSO solvent correction was performed following the Biacore Laboratory guideline 29-0057-18 (GE Healthcare Life Sciences, Buckinghamshire, UK). Drp1i2 was injected at 30 μL/minute for 60 seconds followed by 400 seconds dissociation time, across the chip in a 2-fold dose-response manner from a highest concentration of 200 µM. SPR results were analysed using the Biacore T200 Evaluation Software v.1.

**Mitochondrial morphology analysis**

Human foreskin fibroblasts and Drp1 wildtype and knockout MEF, kindly provided by Hiromi Sesaki (Johns Hopkins University, USA) (2), were transduced with pAS2.EYFP.puro lentiviruses (RNAiCore, Academia Sinica, Taiwan) carrying the mitochondrial-targeted EGFP sequence as described previously (3). Cells were then treated with DRP1i2 (5, 10 or 50 µM) or 0.1% DMSO vehicle control for 24 hours. Images were acquired using Olympus BX60 fluorescence microscope at 600 times magnification and analysed using Fiji imaging software. Cells were categorised into predominantly (>50%) having fused (networked) or fragmented (tubular) mitochondria. At least 100 cells were analysed per treatment group per replicate.

**Drp1 GTPase activity**

The GTPase activity of Drp1 was assessed using a colorimetric GTPase assay kit (Innova Biosciences, Cambridge, UK) according to the manufacturer’s instructions. Test compounds were diluted in DMSO (all with a final concentration of 0.1% DMSO). Recombinant human Drp1 protein isoform-3 (50 ng, final concentration 0.5 µM) was diluted in assay buffer containing 50 mM Tris HCl (pH 7.5) and 2.5 mM MgCl2, then incubated with the test compounds for 15 minutes at 37ºC in a 96 well plate. GTP (0.5 mM, prepared in assay buffer) was subsequently added, and the reaction was incubated for an additional 15 minutes at 37ºC. Drp1-mediated GTP hydrolysis generates GDP and free inorganic phosphate (Pi). Thus, GTPase inhibition was quantified by measuring the amount of Pi released into the assay buffer using a malachite green-based detection dye. The absorbance of the Pi–dye complex was measured at 635 nm using a microplate reader (BenchMark Plus, MS, USA). Values for each experimental condition were normalized to the vehicle (DMSO) control.

**Animal experiments**

### *In vivo model of chronic doxorubicin-induced cardiotoxicity*

To induce chronic doxorubicin-induced cardiotoxicity, Male C57BL/6J mice (male, six weeks old) were given weekly intraperitoneal injections of doxorubicin at 5 mg/kg for five consecutive weeks, resulting in a cumulative dose of 25 mg/kg (4).

*Animal housing and welfare*

Animals were maintained on a chow diet (9% total digestible energy from lipids, Barastoc Irradiated Mice Cubes, Riley, Australia) in a pathogen-free environment under 12-hour light/dark cycle. Animals were group-housed where space permitted to promote normal social behaviour and reduce stress. Animals exhibiting aggression were singly housed to prevent injury. Environmental enrichment, including plastic tubes for hiding and exploration, was provided in all cages.

*Sample size and statistical considerations*

A priori sample size calculations were performed to ensure adequate statistical power to detect a ≥15% improvement in left ventricular ejection fraction, the primary outcome. Based on an expected standard deviation of 10%, a power of ≥0.8, and a significance level (α = 0.05) using one-way ANOVA, a minimum of six animals per group were required.

*Randomisation and blinding*

To minimise potential confounders, animals were randomly assigned to experimental groups using a manual randomisation process. Allocation was not influenced by animal characteristics or investigator preference. The order of treatments and measurements was varied to reduce systematic bias. All animals were maintained under identical environmental conditions, and cages were evenly distributed across racks to minimise positional effects (e.g., differences in temperature, lighting, or airflow). Experimenters performing data collection and analysis were blinded to group allocation.

*Inclusion and exclusion criteria*

All animals that completed the 5-week treatment regimen were included in the final analysis. No animals, experimental units, or data points were excluded post-randomisation. Thus, only complete datasets were included to ensure reliability and comparability of outcomes.

**Echocardiography**

Transthoracic echocardiography was conducted at baseline (day 0) and on day 7, 14, 21, 28 and 35 under light anaesthesia with 2-3% isoflurane in oxygen to evaluate changes in left ventricular function. Imaging was conducted using a Vivid E9 ultrasound system equipped with a 12 MHz phased array probe (GE HealthCare, IL, USA). Two-dimensional parasternal short-axis views were acquired at the level of the papillary muscles, and M-mode tracings were analysed offline using EchoPAC software (GE HealthCare) to measure left ventricular dimension and ejection fraction. After image acquisition, mice were allowed to recover on a heated pad before returning to their cages (total time under anaesthesia was around 15 minutes).

**Cardiac histomorphometry**

Following echocardiographic assessment on day 35, hearts were excised from anaesthetised animals. A mid-ventricular section of the left ventricle was isolated, fixed overnight in 10% neutral buffered formalin at room temperature, and subsequently processed and paraffin-embedded for histological analyses.

*Picrosirius red*

Tissue sections were stained with Picrosirius Red Solution (Abcam, MA, USA) for 20 minutes at room temperature. After incubation, the slides were washed with 0.5% acetic acid (Bio-Strategy, Victoria, Australia) (2 x 1 minute) and rinsed with fresh 100% ethanol (1 x 1 minute followed by 2 x 3 minutes) and cleared in histolene (2 x 2 minutes). Coverslips were mounted onto glass slide using Entellan (Merck Millipore, MA, USA). Collagen fibre-stained sections were captured with BX61 microscope (Olympus). Five random high-power fields (400x magnification) from the left ventricular mid-myocardium were imaged for each sample. The fibrotic area was measured as the percentage of myocardial interstitial fibrosis over total image area in pixels using Fiji software.

*Wheat germ agglutinin*

Tissue sections were stained with rhodamine-conjugated wheat germ agglutinin (50 μg/mL, Vector Labs, CA, USA) for two hours at room temperature in the dark. After incubation, the slides were washed with PBS, coverslipped using Dako fluorescent mounting medium, and stored at 4℃ in dark. The mean cardiomyocyte cross-sectional area was measured in the endomyocardial region. Cell membranes were delineated, and 150 cardiomyocytes oriented perpendicular to the plane of section were analysed. Measurements were obtained from five randomly selected endocardial images acquired at 400× magnification using a BX-61 microscope.

*Griffonia Simplicifolia lectin*

The myocardial vascular density including the capillaries was assessed in tissue sections stained with Griffonia Simplicifolia lectin I. Following antigen retrieval with Proteinase K (Dako, CA, USA) for 6 minutes at room temperature, endogenous peroxidase activity was quenched by incubating sections with 3% hydrogen peroxide in methanol-water mixture (1:1 ratio) for 10 minutes at room temperature. Sections were then incubated with biotin-conjugate Griffonia Simplicifolia lectin (10 μg/mL diluted in Dako antibody diluent, Vector Labs) overnight at 4℃ in a humidified container. The sections were then incubated with conjugated Streptavidin-horseradish peroxidase (2.5 μg/mL diluted in PBS, Dako) for 60 minutes at room temperature, followed by 3,3’-diaminobenzidine substrate (Dako) for 5 minutes. Nuclei were counterstained with Mayer’s haematoxylin. Five random fields from the left ventricular endomyocardium were imaged at 400x magnification for each sample using a BX61 microscope. Lectin-positive vascular structures were counted in Fiji software and vascular density was expressed as the average number of vessels per high-power field and area (mm^2^).

### Proteomics

*Proteomic sample preparation*

Mouse heart tissues were extracted and lysed on ice in lysis buffer (8M urea in 50 mM HEPES pH 8.0) with Halt™ Protease/Phosphatase Inhibitor Cocktail (#78442, Thermo Fisher Scientific) by pulse tip-probe sonication. Protein concentration was quantified with the micro-BCA assay (#23235, Life Technologies, CA, USA). Lysates (10 µg) were processed using the Sera-Mag bead (SP3) workflow as described (5). Briefly, proteins were normalized (10 µg) to a working volume of 50 µL in SDS lysis buffer (1% SDS, 50mM HEPES, pH 8.0) followed by incubation for 5 min at 95°C. Samples were reduced with 10 mM dithiothreitol for 1 hour at 25°C and alkylated with 20 mM iodoacetamide for 30 minutes at 25°C in the dark. Samples were quenched with additional 10 mM dithiothreitol and mixed with carboxylate Sera-Mag Speed Beads mixture (Cytiva, MA, USA) at a 10:1 beads-to-protein ratio (5). Ethanol was added to each sample to a final concentration of 50% (v/v). Proteins were bound to beads by shaking for 10 minutes at 1000 rpm at 25°C and washed three times with 80% ethanol (v/v).

Bead-bound proteins were reconstituted in 50 mM triethylamonium bicarbonate (pH 8.0) with sequence-grade trypsin (V5113, Promega) and Lys-C (121-05063, FUJIFILM Wako Pure Chemical Corporation) at 1:50 and 1:100 enzyme-to-substrate ratios, respectively as detailed (6). Protein digestion was performed overnight at 37°C with agitation at 1,000 rpm. Peptide digests were collected and acidified to a final concentration of 2% (v/v) formic acid. Peptide digests were dried under vacuum centrifugation and reconstituted in 0.07% (v/v) trifluoroacetic acid in MS-grade water. Subsequently, peptides were quantified by Fluorometric Peptide Assay (#23290, Thermo Fisher Scientific) as per manufacturer’s instructions.

*Liquid Chromatography-Mass Spectrometry (LC-MS) analysis*

Peptide samples were analysed using a Thermo Scientific Vanquish Neo UHPLC system in trap-and-elute mode coupled to an Orbitrap Astral mass spectrometer with an Easy-spray ion source. Peptides (50 ng) were loaded onto a PepMap™ Neo Trap Cartridge (#174500, Thermo Fisher Scientific) and separated on an EASY-Spray PepMap nanoflow 150 mm RSLC C18 UHPLC column with an integrated emitter (#ES906; Thermo Fisher Scientific) maintained at 55°C. Mobile phase A consisted of 0.1% formic acid in water, and mobile phase B consisted of 80% acetonitrile with 0.1% formic acid. Peptides were eluted using a 13.5-min gradient at 1.8 µL/min: 4% B for 0.7 min, increasing to 8% B at 1.0 min, 28% B at 9.7 min, 50% B at 13.2 min, and 99% B at 13.7 min, followed by a 0.7-min wash at 99% B at 2.5 µL/min.

Full MS scans were acquired in the Orbitrap at 240,000 resolution, over an m/z range of 380–980 with an AGC target of 5 × 10⁶, maximum injection time of 5 ms and 1 microscan. MS/MS spectra were collected in the Astral analyser using 200 isolation windows of 3 m/z width with 0 m/z overlap with the same precursor range, AGC 5 × 10⁴, maximum injection time of 5 ms and 1 microscan. Fragmentation was performed using higher-energy collisional dissociation with normalized collision energy of 25%, and fragment ions were detected from m/z 150–2000. The total cycle time was approximately 0.6 s. Data was acquired using Xcalibur software v4.7 SP1 (Thermo Fisher Scientific). Mass spectrometry-based proteomics and phospho-proteomics data is deposited to the ProteomeXchange Consortium via the MASSive partner repository and available via MASSive with identifier (proteome: MSV000099244).

*Data processing and analysis*

All raw data were analysed using DIA-NN neural network and interference correction (v2.0) in library-free mode searched against UP000000589_MOUSE (Nov 2024) reference proteome (54742 entries) (7) Spectral libraries were predicted using the deep learning algorithm employed in DIA-NN, generated from the mouse reference proteome with Trypsin/P, allowing up to 1 missed cleavage.

The DIA-NN search included the following settings: Protein inference = ‘Genes’, Neural network classifier = ‘Single-pass mode’, Quantification strategy = ‘Robust LC (high precision)’, Cross-run normalization = ‘RT-dependent’, Library Generation = ‘IDs, RT and IM Profiling’ and Speed and RAM usage = ‘Optimal results’. Mass accuracy and MS1 accuracy were set to 0 for automatic inference. ‘No share spectra’, ‘Heuristic protein inference’ and ‘MBR’ were checked. Mass spectra were analysed with false discovery rate of 1% for precursor identifications. The precursor charge range was 1-4, and the m/z precursor range was set to 300-1800 for peptides consisting of 7-30 amino acids with N-term methionine excision and cysteine carbamidomethylation enabled as a fixed modification with 0 maximum number of variable modifications. Downstream data analysis was performed using Perseus (v2.0.11.0) of the MaxQuant software package with bar plots/violin plots generated using GraphPad Prism (version 10) or Microsoft Excel. Stringent data quality inclusion was further applied with minimum 3 protein group quantifications in at least one group. For samples that did not meet protein identification depth/coverage or due to ultra-low peptide recovery, data exclusions were applied (with minimum N=4 independent biological replicates per group applied). Protein intensities were log2 transformed and normalized using quantile normalization. Principal component analysis (PCA) was conducted using features with valid measurements in at least 70% of samples within at least one group. Log2‑transformed intensity values from all samples were used, and missing values were imputed from a normal distribution (downshift = 1.8, width = 0.3; Perseus). Imputed values were used exclusively for PCA and were omitted from all subsequent statistical analyses. For differential analysis, proteins were subjected to ANOVA analysis and Student’s t-test, with hierarchical clustering performed using Euclidian distance and average linkage clustering. MitoCarta 3.0 were used for mitochondrial annotation and tissue association analyses (8). Gene Ontology 1D-enrichment analysis was performed in Perseus based on log2 fold changes for sample groups (P < 0.05). Data visualization was performed using Perseus, R (ggplot2) package.

**Human iPSC culture**

Human iPSC-Foreskin-2 cell line, generously provided by James A. Thomson (University of Wisconsin) (9) was maintained on vitronectin-coated plates (Thermo Fisher Scientific, MA, USA) in TeSR-E8 medium (Stem Cell Technologies, VA, Canada). The culture medium was refreshed every 2-3 days, and cells were passaged on a weekly basis using ReLeSR (Stem Cell Technologies).

**Human iPSC cardiomyocyte differentiation**

Human iPSCs were differentiated into cardiomyocytes using an established protocol (10). Briefly, iPSCs were dissociated into single cells and plated on human embryonic stem cell (hESC)-qualified Matrigel basement membrane matrix (Corning, NY, USA) coated plates at a density of 1.25 x 10^5^ cells/cm² in TeSR-E8 medium supplemented with 10 μM Y-27632 dihydrochloride (Tocris Bioscience, Bristol, UK), referred to as day -2. On day 0, the medium was changed to RPMI-1640 medium containing 1x B27 without insulin supplement (Thermo Fisher Scientific), growth factor reduced (GFR) Matrigel (1:60 dilution) (Corning), and CHIR99021 (10 μM, Cayman Chemical, MI, USA) for 24 hours. On day 1, the medium was changed to RPMI-1640 basal medium containing 1x B27 without insulin supplement for 24 hours. On day 2, the medium was replaced with RPMI-1640 basal medium containing 1x B27 without insulin supplement and 5 μM IWP2 (Sigma-Aldrich, MA, USA) for 72 hours. From days 5 to 12, cells were cultured in RPMI-1640 medium containing 1x B27 supplement and 20 ng/mL ascorbic acid 2-phosphate sesquimagnesium salt hydrate (Sigma-Aldrich) (cardiomyocyte medium), and medium was refreshed every 48-72 hours. On day 12, the cells were dissociated into single cells and replated on hESC-qualified Matrigel-coated plates at a ~1:4 ratio and cultured in DMEM/F-12 + GlutaMAX medium supplemented with 20% foetal calf serum, 100 μM β-mercaptoethanol, 100 μM non-essential amino acids, 50 U/mL penicillin/streptomycin, and 10 μM Y-27632 dihydrochloride. On day 13, the medium was replaced with cardiomyocyte medium. From days 14-19, cardiomyocytes were enriched in DMEM (without glucose) medium (Thermo Fisher Scientific) supplemented with 4 mM lactate (Sigma-Aldrich). On day 19, enriched cardiomyocytes were dissociated and plated on hESC-qualified Matrigel-coated 48-well plates at a cell density of 2.5 x 10^4^ cells/cm^2^ in DMEM/F-12 GlutaMAX medium supplemented with 20% foetal calf serum, 100 μM mecaptoethanol, 100 μM non-essential amino acids, 50 U/mL penicillin/streptomycin, and 10 μM Y-27632 dihydrochloride. From day 20 onwards, cardiomyocytes were cultured in cardiomyocyte medium, and medium was replaced every 2-3 days. Purity of cardiomyocytes was assessed by immunocytochemistry to quantify the percentage of cells expressing the cardiac-specific marker cardiac troponin-T. A minimum purity level of 95% for cardiomyocytes was required to proceed with experiments. Cells were replated at a density of 2.5 x 10^4^ cells/ well (7.81 x 10^4^ cells/ cm^2^) in a 96-well flat bottom plates for 2D monoculture experiments or were used to generate 3D microtissues. Cells were treated for 3 days with vehicle control (0.1% DMSO), DRP1i2, doxorubicin, or a combination of doxorubicin and DRP1i2.

**Human iPSC cardiac fibroblast differentiation**

To generate cardiac fibroblasts, human iPSCs were first differentiated into epicardial cells, as described previously with modifications (10,11). Briefly, iPSCs were plated on hESC-qualified Matrigel-coated plates at a density of 2.5 x 10^4^ cells/cm^2^ in TeSR-E8 medium supplemented with 10 μM Y-27632 dihydrochloride, referred to as day -1. On day 0, cardiac mesoderm was induced by replacing TESR-E8 medium with APEL medium (Stem Cell Technologies) supplemented with 20 ng/mL BMP4 (R&D Systems, MN, USA), 20 ng/mL activin A (Peprotech, NJ, USA) and 1.5 μM CHIR99021. On day 3, cytokines were removed and replaced with APEL medium supplemented with 5 μM IWP2, 30 ng/mL BMP4 and 1 μM retinoic acid (Sigma Aldrich) for 3 days. On day 6, medium was replaced with APEL medium supplemented with 30 ng/mL BMP4 and 1 μM retinoic acid. On day 9, cells were dissociated and plated on human fibronectin-coated plates (Merck Millipore, MA, USA) at a density of 1.5 x 10^4^ cells/cm^2^ in APEL medium supplemented with 10 μM SB431542 (Tocris Bioscience) and 10 μM Y-27632 dihydrochloride. On day 13, cardiac fibroblast differentiation was induced using an established protocol (11). Briefly, confluent epicardial cells were plated on vitronectin-coated plates at a density of 2.5 x 10^4^ cells/cm^2^ in APEL medium supplemented with 10 ng/mL FGF2 (R&D Systems) and 10 μM Y-27632 dihydrochloride for 2 days. On day 15 and 17, medium was refreshed with APEL medium supplemented with 10 ng/mL FGF2. From day 20 to 29, cardiac fibroblasts were expanded in Fibroblast Growth Medium 3 (FGM3; PromoCell, Heidelberg, Germany), and medium was replaced every 2-3 days. Cells were replated at a density of 2.5 x 10^4^ cells/well (7.81 x 10^4^ cells/cm^2^) in a 96-well flat bottom plates for 2D monoculture experiments or were used to generate 3D microtissues. Cells were treated for 3 days with vehicle control (0.1% DMSO), DRP1i2, doxorubicin, or a combination of doxorubicin and DRP1i2.

**Human iPSC endothelial cell differentiation**

Human iPSCs were differentiated into CD31^+^ endothelial cells, following an established protocol (12,13). Briefly, iPSCs were first dissociated into single cells and plated on hESC-qualified Matrigel-coated plates at a density of 1 x 10^5^ cells/cm^2^ in TESR-E8 media supplemented with 10 μM Y-27632 dihydrochloride. After 24 hours, which is referred to as day 0, the culture medium was replaced with DMEM/F12 + GlutaMAX medium containing 1x N-2 supplement (Thermo Fisher Scientific), 1x B27 supplement, 8 μM CHIR99021, and 25 ng/mL BMP4 (Stem Cell Technologies) for 3 days. On day 3, the culture medium was replaced with StemPro-34 SFM complete medium (Thermo Fisher Scientific) supplemented with 200 ng/mL of VEGF-165 (Peprotech) and 2 μM forskolin (Sigma-Aldrich) for 3 days. On day 6, endothelial cells were dissociated into single-cell suspensions, incubated with a FITC-conjugated mouse anti-human CD31 antibody (20 μg/mL; clone WM59; #555445, BD Biosciences, NJ, USA), and sorted using a BD FACS Aria III cell sorter (BD Biosciences). CD31⁺ cells were subsequently expanded on fibronectin-coated plates and cultured in EGM2-MV medium (Lonza, Basel, Switzerland) supplemented with 50 ng/mL VEGF-165. Cells were replated at a density of 2.5 x 10^4^ cells/well in a 96-well flat bottom plates for 2D monoculture experiments or were used to generate 3D microtissues. Cells were treated for 3 days with vehicle control (0.1% DMSO), DRP1i2, doxorubicin, or a combination of doxorubicin and DRP1i2.

**Human iPSC vascular smooth muscle cell differentiation**

Human iPSCs were differentiated into vascular smooth muscle cells, using established protocols with modifications (10,14). Briefly, iPSCs were dissociated and plated on GFR Matrigel-coated plates at a density of 4 x 10^4^ cells/cm^2^ in TeSR-E8 medium supplemented with 10 μM Y-27632 dihydrochloride. After 24 hours, referred to as day 0, medium was replaced with N2/B27 medium consisting of a 1:1 ratio of DMEM/F12 + GlutaMAX medium and neurobasal medium (Thermo Fisher Scientific), supplemented with 1x N2 supplement (Thermo Fisher Scientific), 1x B27-Vitamin A supplement (Thermo Fisher Scientific), 8 μM CHIR99021 and 25 ng/mL BMP4 (Stem Cell Technologies). On day 3, the medium was replaced with N2/B27 medium supplemented with 10 ng/mL PDGF-BB (Peprotech) and 2 ng/mL activin A (Peprotech). On day 4, cells were replated on collagen-coated plates at a density of 8 x 10^5^ cells/cm^2^ in N2/B27 medium supplemented with 2 μg/mL heparin (Stem Cell technologies) and 2 ng/mL activin A. On day 7, medium was replaced with SmGM-2 medium (Lonza). Cells were replated at a density of 2.5 x 10^4^ cells/well in a 96-well flat bottom plates for 2D monoculture experiments or were used to generate 3D microtissues. Cells were treated for 3 days with vehicle control (0.1% DMSO), DRP1i2, doxorubicin, or a combination of doxorubicin and DRP1i2.

### Cardiac microtissue

*Cardiomyocyte-only microtissue*

Cardiomyocyte-only microtissues were generated using previously published protocols with minor modifications (15). Briefly, human iPSC-derived cardiomyocytes (day 19) were seeded into 96-well round bottom ultra-low attachment plate at 7.5 x 10^4^ cells/well in 20 μL of DMEM/F-12 GlutaMAX medium supplemented with 20% foetal bovine serum, 0.1 mM β-mercaptoethanol, 0.1 mM nonessential amino acids, 50 U/mL penicillin/streptomycin and 10 μM Y-27632 dihydrochloride. After 2 hours of incubation at 37 °C in a humidified atmosphere of 5% CO₂/95% air, 80 μL of cardiomyocyte medium (RPMI-1640 supplemented with 1× B27 (Thermo Fisher Scientific) and 20 ng/mL ascorbic acid 2-phosphate sesquimagnesium salt hydrate (Sigma-Aldrich)) supplemented with 20% methylcellulose (Sigma Aldrich) was added to facilitate microtissue compaction. After 24 hours, the medium was replaced with cardiomyocyte medium. Microtissues were treated for 3 days with vehicle control (0.1% DMSO), DRP1i2 (10, 50, or 100 μM), doxorubicin, or a combination of doxorubicin and DRP1i2.

### *Multicellular cardiac microtissue*

Multicellular cardiac microtissues were constructed using previously published protocols with modifications (10,15). The multicellular cardiac microtissues were assembled with the same total cell number as cardiomyocyte-only microtissues (7.5 x 10^4^ cells/microtissue) in 96-well round bottom ultra-low attachment plate. Each microtissue is composed of 75% cardiomyocytes (5.625 x 10^4^ cells/well), 20% endothelial cells (1.5 x 10^4^ cells/well), 2.5% cardiac fibroblasts (1.875 x 10^3^ cells/well) and 2.5% vascular smooth muscle cells (1.875 x 10^3^ cells/well) per well. Cells were resuspended in 20 μL of a mixture of cardiomyocyte medium, EGM2-MV medium (Lonza, Basel, Switzerland), SmGM-2 (Lonza) and FGM-3 (PromoCell, Heidelberg, Germany) at a 1:1:1:1 ratio (referred to as multicellular medium), supplemented with 10 μM Y-27632 dihydrochloride. After 2 hours of incubation at 37 °C in a humidified atmosphere of 5% CO₂/95% air, 80 μL of multicellular medium containing 20% methylcellulose was added to facilitate microtissue compaction. After 24 hours, the medium was replaced with multicellular medium. Microtissues were treated for 3 days with vehicle control (0.1% DMSO), DRP1i2 (10, 50, or 100 μM), doxorubicin, or a combination of doxorubicin and DRP1i2.

**Human cancer cell culture**

MDA-MB-231 breast cancer cells and A549 lung cancer cells were cultured in Dulbecco's Modified Eagle's Medium (Sigma-Aldrich) supplemented with 2 mM GlutaMAX (Thermo Fisher Scientific) and 10% foetal bovine serum (Sigma-Aldrich). OVCAR3 ovarian cancer cells were cultured in RPMI-1640 medium supplemented with 2 mM GlutaMAX and 10% foetal bovine serum. MG63 osteosarcoma cells were cultured in alpha-Modified Eagle’s Medium (Thermo Fisher Scientific) supplemented with 2 mM GlutaMAX and 10% foetal bovine serum. All the cancer cells were maintained in a humidified 5% CO_2_/95% air incubator at 37°C and medium was changed every 2-3 days. 2D cancer cells were treated for 5 days with vehicle control (0.1% DMSO), DRP1i2, doxorubicin, or a combination of doxorubicin and DRP1i2.

To generate cancer spheroids, MG63 osteosarcoma cells were seeded into 96-well ultra-low attachment round bottom plates (Corning, NY, USA) at 2.0 x 10^4^ cells/well in 20 μL of culture medium. After incubation at 37°C for two hours, 80 μL of culture medium supplemented with 20% methylcellulose was added to cancer cells to promote spheroid compaction. After 24 hours, the medium was replaced with culture medium without methylcellulose and maintained in a humidified 5% CO_2_/95% air incubator at 37°C with medium changed every 2-3 days. MG63 osteosarcoma spheroids were treated for 5 days with vehicle control (0.1% DMSO), DRP1i2 (50 μM), doxorubicin (0.05 μM), or a combination of doxorubicin and DRP1i2.

BJ1 fibroblasts were cultured in DMEM/F-12 medium supplemented with 10% foetal bovine serum and 100 U/mL penicillin/streptomycin, while FT282 cells, a non-transformed epithelial cell line derived from the fallopian tube, were maintained in DMEM/F-12 medium supplemented with 10% knockout FBS. Non-cancerous human ovarian surface epithelial cells (HOSE 6.3 and HOSE 17) were cultured in RPMI-1640 medium containing 2 mM GlutaMAX and supplemented with 10% foetal bovine serum.

**Video contractility**

Bright field videos of contracting 2D cardiomyocytes and 3D cardiac microtissues were acquired at 200x and 40x magnifications, respectively, at 60 frames/second using an inverted IX71 microscope (Olympus) equipped with a DP74 digital camera (Olympus). Beating cardiomyocytes or cardiac microtissues were analysed using the MUSCLEMOTION plugin in ImageJ (16) to extract contraction kinetics, including contraction duration, time to peak, time to relax, and beat rate variability. Beat rate variability was quantified as the root mean square of successive differences (RMSSD) normalized to the peak-to-peak interval (RMSSD/RR). RMSSD was calculated as the square root of the mean of the squared differences between consecutive peak-to-peak (RR) intervals (12).

**Mitochondrial ROS production**

The level of mitochondrial ROS was assessed using MitoSOX Red reagent (Thermo Fisher Scientific). Briefly, 2D cells or 3D cardiac microtissues were stained with 5 μM MitoSOX Red diluted in HBSS+/+ and incubated in a humidified 5% CO_2_/95% air incubator at 37°C for 20 minutes. 2D cancer cells were imaged at 100x magnification using the Operetta High Content Imaging System (PerkinElmer, MA, USA). Three fields of view per well was captured under appropriate filter settings. Automated segmentation in Harmony software identified the nuclei and corresponding cytoplasmic regions of interest, allowing quantification of fluorescent intensity on a per-cell basis. Background subtraction was applied to reduced non-specific signal, and the mean fluorescence intensity was averaged across all fields within a well to obtain normalised values for analysis. 2D cardiac cells were imaged at 200x magnification using an IX71 microscope (Olympus). At least 100 cells from 3-5 random fields in each sample were analysed. 3D cardiac microtissues were imaged at 50x magnification using the Thunder microscope (LEICA Microsystems, Wetzlar, Germany). The MitoSOX Red fluorescence intensity was analysed with ImageJ software and calculated using the corrected total cell fluorescence method (17).

Total cellular ROS was measured using CellROX Green reagent. 3D cancer spheroids were stained with 5 μM CellROX Green reagent diluted in HBSS+/+ and incubated in a humidified 5% CO_2_/95% air incubator at 37°C for 20 minutes. After incubation, cells were washed twice with HBSS+/+ before replacing with fresh HBSS+/+ prior to imaging. 3D spheroids were imaged at 50x magnification using the Thunder microscope (LEICA Microsystems). The CellROX Green reagent fluorescence intensity was analysed with ImageJ software and calculated using the corrected total cell fluorescence method.

**Mitochondrial membrane potential**

Mitochondrial membrane potential was assessed using tetramethylrhodamine methyl ester (TMRM, Thermo Fisher Scientific). 2D cells were stained with 10 nM TMRM diluted in HBSS+/+ and incubated in a humidified 5% CO2/95% air incubator at 37°C for 20 minutes. 2D cancer cells were imaged at 100x magnification using the Operetta High Content Imaging System. Three fields of view per well was captured under appropriate filter settings. Automated segmentation in Harmony software identified the nuclei and corresponding cytoplasmic regions of interest, allowing quantification of fluorescent intensity on a per-cell basis. Background subtraction was applied to reduced non-specific signal, and the mean fluorescence intensity was averaged across all fields within a well to obtain normalised values for analysis. 2D cardiac cells were imaged at 200x magnification using an IX71 microscope (Olympus). At least 100 cells from 3-5 random fields in each sample were analysed. 3D spheroids were imaged at 50x magnification using the Thunder microscope (Leica). The TMRM fluorescence intensity was quantified using ImageJ software and calculated using the corrected total cell fluorescence method.

**Immunocytochemistry**

2D cells and 3D spheroids were fixed with 10% neutral buffered formalin (Sigma-Aldrich) at room temperature for 20 minutes and 1 hour respectively. 3D cardiac microtissues were then dehydrated in 20% sucrose solution for 24 hours. The dehydrated samples were embedded in Optimal Cutting Temperature compound and cryosectioned into 5 μm thick sections. Sections were treated with 0.2% Triton X-100 permeabilisation buffer for 10 minutes followed by Ultra V Block solution for another 10 minutes. The sections were then stained with primary antibodies; anti-cardiac troponin T (2 μg/mL, rabbit polyclonal, Abcam), anti-cardiac troponin T (4 μg/mL, mouse monoclonal, clone 1C11, Abcam), anti-alpha actinin (25 µg/mL, mouse monoclonal, A7811, Sigma-Aldrich), anti-CD31 (4 μg/mL, mouse monoclonal, clone JC70A, Dako), anti-vimentin (0.3 μg/mL, mouse monoclonal, clone V9, Dako), or anti-SM22-⍺ (5 μg/mL, rabbit polyclonal, Abcam) at 4℃ overnight, followed by Alexa Fluor-488-conjugated goat-anti-rabbit or goat-anti-mouse (10 μg/mL, Invitrogen) and Alexa Fluor-594-conjugated goat-anti-mouse or goat-anti-rabbit (10 μg/mL, Invitrogen) secondary antibodies. Sections were then counterstained with 1 μg/mL of DAPI (Invitrogen) for nuclear staining. Epifluorescence images of immunostained sections were acquired with a BX61 microscope (Olympus) using analySIS software.

Mitochondrial morphology in 2D MG63 osteosarcoma cells was assessed by immunofluorescent staining for Hsp60. Cells were incubated overnight at 4°C with anti-Hsp60 antibody (1.58 μg/mL, rabbit polyclonal IgG, Abcam, ab46798), followed by Alexa Fluor 488 goat anti-rabbit IgG (10 μg/mL, Thermo Fisher Scientific, A11008) and nuclear counterstaining with DAPI (1 μg/mL) for 1 hour at room temperature. Images were acquired using an upright Olympus BX61 microscope at 600× magnification. Images were acquired using an upright Olympus BX61 microscope at 600× magnification. At least 60 cells were analysed from 10-15 randomly acquired images per group per replicate.

**Cell viability**

The viability of 2D cells was assessed using the CellTiter-Blue assay (Promega, WI, USA). Briefly, culture media was removed and replaced with 60 μL diluted dye consists of 50 μL of cell culture media + 10 μL of dye. The plate was then incubated for 4 hours in a humidified 5% CO2/95% air incubator at 37°C. The fluorescent signal was measured at 560/590 nm (excitation and emission peaks of resorufin) using the EnSpire Multimode Plate Reader (PerkinElmer). All readings were corrected for the background signals (MTT reagent + serum-free medium without cells), and the fold change of cell viability was expressed over the vehicle control.

The viability of 3D cancer spheroids was assessed using the CellTiter-Glo assay (Promega). Briefly, culture media was removed and replaced with 100 μL of mixed dye consists of 50 μL of cell culture media + 50 μL of dye. The plate was then vigorously shaken for 5 minutes to induce cell lysis. After that, the plate was incubated at room temperature for 20 minutes to stabilise the luminescent signal, which was then recorded using the PolySTAR plate reader (BMG Labtech). All readings were corrected for the background signals (CellTiter-Glo reagent + cell culture media with no cells) and the fold change in cell viability was expressed over the control.

**Cell metabolism**

The Seahorse XFe96 Extracellular Flux Analyzer (Agilent Technologies, CA, USA) was used to measure oxygen consumption rate (OCR) and extracellular acidification rate (ECAR) using the Seahorse XF Cell Mito Stress Test and Glycolysis Stress Test, respectively. For 2D MG63 osteosarcoma cells, cells were treated for 5 days with vehicle control, DRP1i2, doxorubicin, or a combination of doxorubicin and DRP1i2. After treatment, cells were seeded at 1.5 × 10^4^ cells/well in a Seahorse XFe96 microplate (5-6 technical replicates) and incubated overnight at 37°C in 5% CO₂ prior to the assay. For cardiac microtissues, microtissues were transferred into Matrigel-coated Seahorse XFe96-Spheroid microplates and incubated for 2 days, followed by treatment for 3 days with vehicle control, DRP1i2, doxorubicin, or a combination of doxorubicin and DRP1i2.

On the day of the Seahorse assay, for the Mito Stress Test, cells were washed and media replaced with OCR assay medium (Seahorse XF Base Medium supplemented with 25 mM glucose, 1 mM sodium pyruvate and 1 mM glutamine, pH 7.4). For the Glycolysis Stress Test, cells were washed and media replaced with ECAR assay medium (Seahorse XF Base Medium supplemented with 1 mM L-glutamine, pH 7.4). The cells were then equilibrated in a non-CO_2_ incubator at 37°C for 30 minutes prior to the assay.

The assay protocols consisted of repeated cycles with simultaneous assessment of OCR and ECAR. For 2D cultures, each cycle comprised 2 minutes of mixing, 30 seconds of waiting, and 3 minutes of measuring; for 3D microtissues, cycles comprised 0 min of mixing (mixing was omitted to prevent microtissue dislodgement), 2 minutes of waiting, and 3 minutes of measuring. For the Mito Stress Test, OCR was measured under basal conditions and after sequential injections of oligomycin (1 μM), carbonyl cyanide p-trifluoromethoxy-phenylhydrazone (FCCP, 1 μM) and rotenone/antimycin A (1 μM each) to assess basal respiration, ATP-linked respiration, proton leak, uncoupled respiration, spare respiratory capacity and non-mitochondrial oxygen consumption. For the Glycolysis Stress Test, ECAR was measured under glucose starvation conditions and after sequential injections of glucose (10 mM), oligomycin (1 μM) and 2-deoxy-D-glucose (2-DG, 50 mM) to assess glycolysis, glycolytic capacity and non-glycolytic acidification. Data were collected with the Seahorse XFe96 Analyzer and analysed using Wave software (Agilent Technologies). OCR and ECAR values were normalised to cell number per well.

**Western blotting**

Proteins were extracted from cells using RIPA lysis buffer (Thermo Fisher Scientific) supplemented with 1x phosphatase and protease inhibitor cocktail (Thermo Fisher Scientific). Protein concentrations were determined with a bicinchoninic acid assay kit (Thermo Fisher Scientific). Proteins were reduced and denatured using lithium dodecyl sulphate sample buffer containing 50 mM dithiothreitol followed by heating at 70ºC for 10 minutes. For electrophoresis, 30 µg of protein was loaded into 12% Bis-Tris protein gel (Thermo Fisher Scientific) and ran at 150 V for 60 minutes. The proteins on the gel were then transferred onto a polyvinylidene difluoride membrane (GE Healthcare Life Sciences, Australia) at 30 V for 60 minutes and blocked with Intercept Tris-buffered saline blocking buffer (LiCOR Biosciences, NE, USA) for 60 minutes at room temperature. The membrane was washed in Tris-buffered saline containing 0.1% Tween-20 (TBS-T) and incubated with primary antibodies at 4ºC overnight. Primary antibodies used include phosphorylated Drp1-Ser616 (1 µg/mL, rabbit monoclonal IgG, Cell Signalling, #3455), phosphorylated Drp1-Ser637 (1 µg/mL, rabbit polyclonal IgG, Cell Signaling, #4867), total Drp1 (0.33 µg/mL, mouse monoclonal IgG, Abcam, ab56788), and β-actin (1 µg/mL, mouse monoclonal AC-15, Abcam, ab8226). Following 3 washes with TBS-T, the membrane was incubated with secondary antibody at room temperature for 60 minutes. Secondary antibodies include Alexa-680 Donkey-anti-rabbit (0.25 µg/mL; Thermo Fisher Scientific, #A10043) and IRDye 800CW Goat anti-mouse (0.25 µg/mL; LiCOR Biosciences, #926-32210). After 3 additional washes, the membrane was scanned at 700 nm and 800 nm wavelengths using the Odyssey CLx scanner (LiCOR Biosciences). Intensity of protein bands was semi-quantified using volume box tools in Image Studio software (Image Studio Lite Ver 5.2).

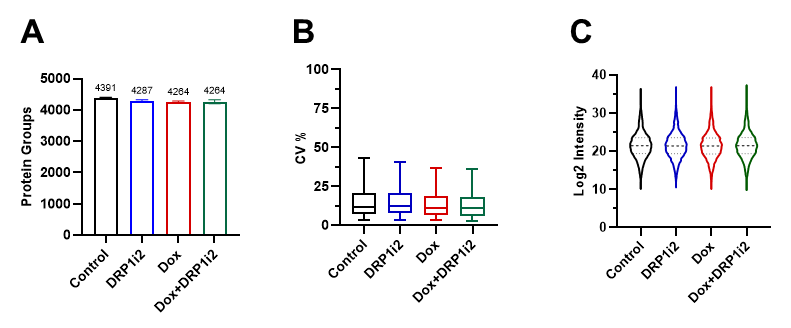

***Supplementary Figure 1. Cardiac proteomics data quality and group distribution.*** *Global proteome profiling of murine heart tissue in response to doxorubicin-induced cardiotoxicity.* ***(A)*** *Average number of protein groups detected in each experimental prior to data filtering. Error bars represent standard error of mean (n=4).* ***(B)*** *Boxplot showing the distribution of coefficient of variation (CV) values for raw protein group intensities within each group, displayed from the 5^th^ to 95^th^ percentile.* ***(C)*** *Violin plot illustrating the distribution of log2-transformed label-free quantification intensities across all samples.*

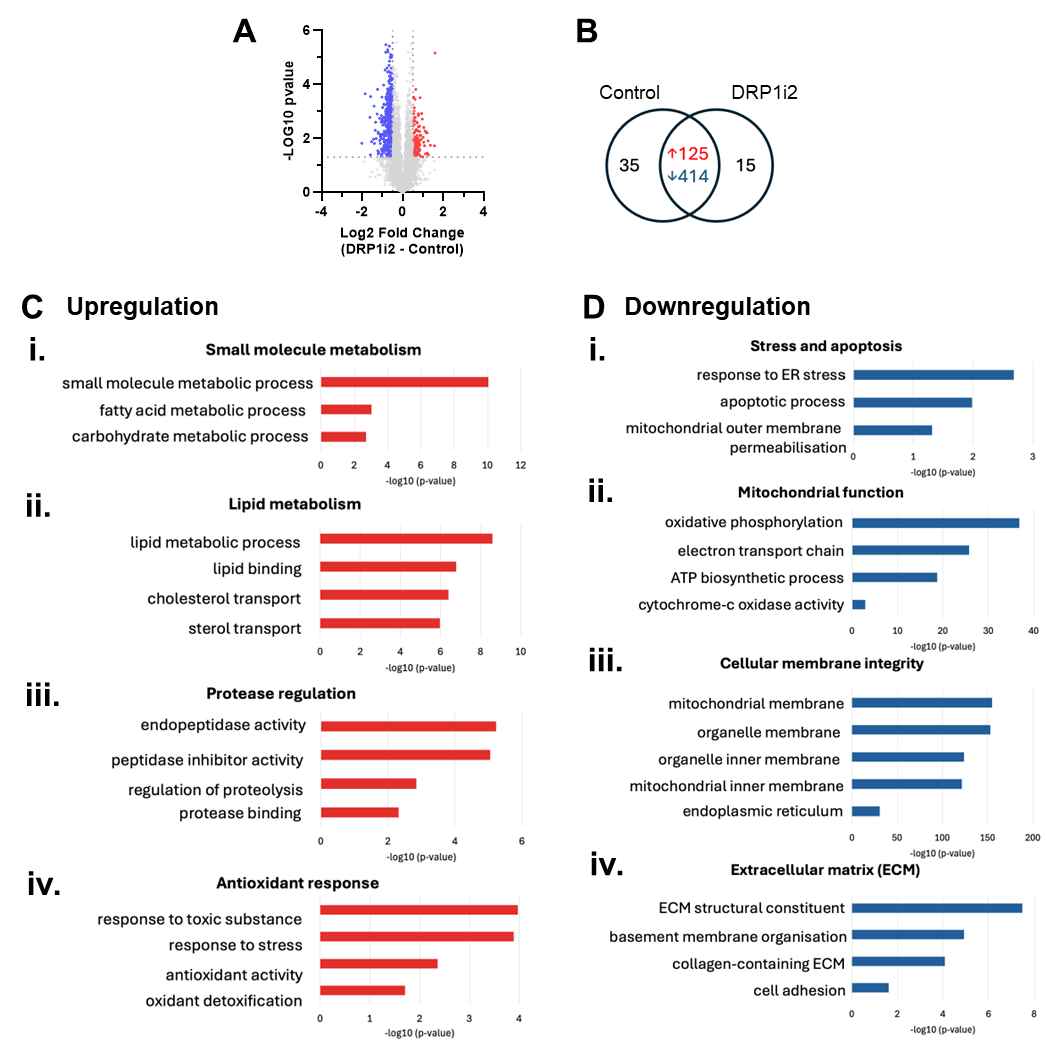

***Supplementary Figure 2. DRP1i2-induced cardiac proteomic remodelling reflects a non-toxic adaptive cellular response****.* ***(A****) Volcano plot showing differential protein expression in DRP1i2-treated hearts versus control, based on Student’s t-test. Significantly downregulated proteins in DRP1i2 group are shown in blue (p < 0.05 and log2 fold change <-0.5), and upregulated proteins are shown in red (p < 0.05 and log2 fold change > 0.5). Data quality inclusion was applied with minimum 3 protein group quantifications in at least one group. (B) Venn diagram illustrating the number of significantly regulated and unique proteins in the DRP1i2 versus control comparison.* ***(C-D)*** *Bar graphs showing gene ontology enrichment analysis using g:Profiler for proteins significantly upregulated and unique to DRP1i2* ***(C)****, and proteins significantly downregulated in OB37 and unique to the vehicle control group* ***(D)****. Enriched pathways are ranked by gene ratio and –log10 p-value. n=4 biological replicates per group.*

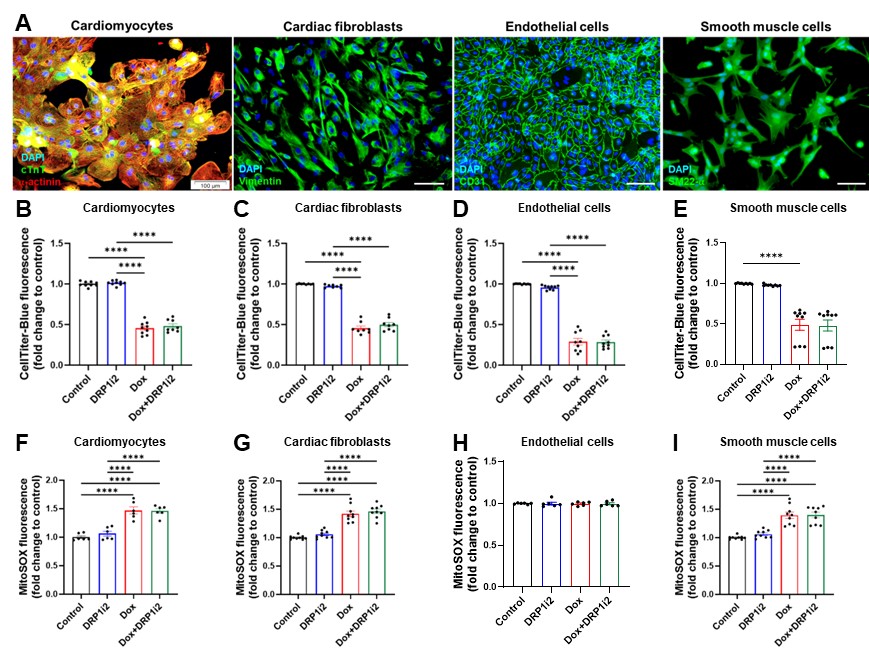

***Supplementary Figure 3. DRP1i2 does not attenuate doxorubicin-induced cytotoxicity in 2D human iPSC-derived cardiac cells. (A)*** *Representative immunofluorescence images of human iPSC-derived cardiac cell types: cardiomyocytes stained for cardiac troponin T (cTnT) and α-cardiac actinin; cardiac fibroblasts stained for vimentin; endothelial cells stained for CD31; and smooth muscle cells stained for SM22-α. Nuclei were counterstained with DAPI. Scale bar = 100 µm.* ***(B-E)*** *Cell viability assessed by CellTiter-Blue assay in cardiomyocytes* ***(B),*** *cardiac fibroblasts* ***(C)****, endothelial cells* ***(D)****, and smooth muscle cells* ***(E)*** *(n = 8-9).* ***(F-I)*** *Representative images and quantification of mitochondrial reactive oxygen species (ROS) production in cardiomyocytes* ***(F)****, cardiac fibroblasts* ***(G)****, endothelial cells* ***(H)****, and smooth muscle cells* ***(I)*** *(n = 6-9). ****P<0.0001 by one-way ANOVA.*

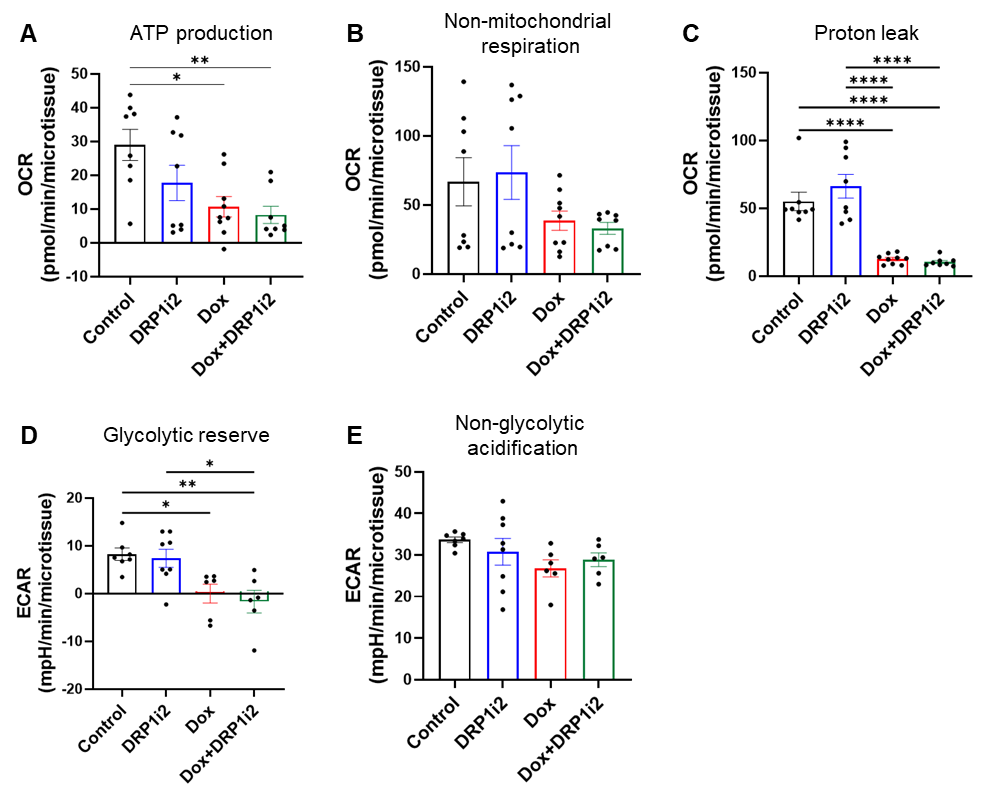

***Supplementary Figure 4. Seahorse extracellular flux analysis of energy metabolism in multicellular cardiac microtissues. (A-C)*** *Mitochondrial respiration assessed by oxygen consumption rate (OCR, n = 8-9 microtissues).* ***(A)*** *ATP-linked respiration.* ***(B)*** *Non-mitochondrial respiration.* ***(C)*** *Proton leak.* ***(D-E)*** *Glycolytic function assessed by extracellular acidification rate (ECAR, n = 6-8 microtissues).* ***(D)*** *Glycolytic reserve.* ***(E)*** *Non-glycolytic acidification.* *Data are presented as mean ± SEM. *P<0.05, **P<0.01, ****P<0.0001 by one-way ANOVA.*

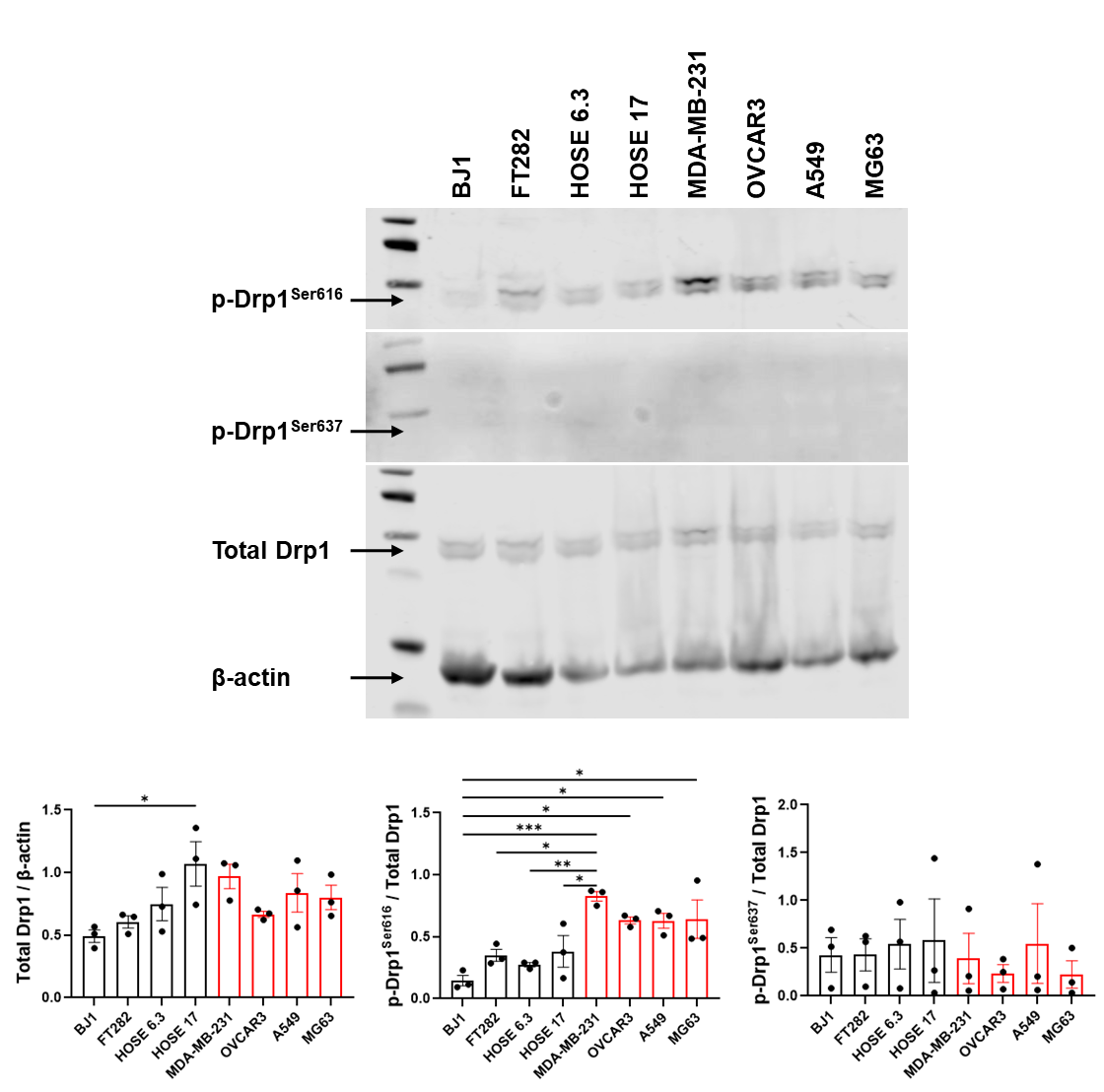

***Supplementary Figure 5. Expression of Drp1 and phosphorylated Drp1 in human cancer cell lines.*** *Representative immunoblots and semi-quantitative analyses of total Drp1, phosphorylated Drp1 at Ser616 (p-Drp1^Ser616^), phosphorylated Drp1 at Ser637 (p-Drp1^Ser637^), and β-actin in control cells (BJ1 foreskin fibroblasts, FT282 fallopian tube epithelial cells and HOSE ovarian surface epithelial cells) and in MDA-MB-231 breast cancer, OVCAR3 ovarian cancer, A549 lung cancer, and MG63 osteosarcoma cell lines. Data are presented as mean ± SEM. *P < 0.05, **P < 0.01, ***P < 0.001, as determined by one-way ANOVA.*

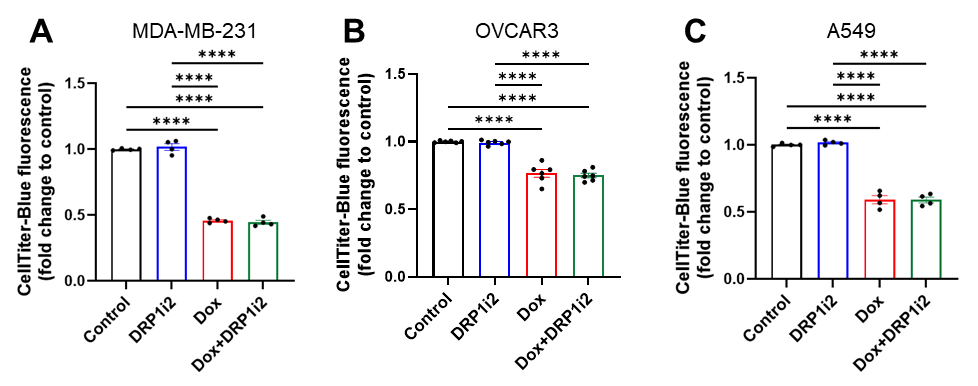

***Supplementary Figure 6. Anti-cancer effects of doxorubicin and DRP1i2 in 2D cancer cell cultures.*** *Cell viability of MDA-MB-231 breast cancer* ***(A)****, OVCAR3 ovarian cancer* ***(B)****, and A549 lung cancer* ***(C****) cells following 5-day treatment with vehicle control (0.1% DMSO), DRP1i2, doxorubicin, or a combination of doxorubicin and DRP1i2 (n=4-6). Data represent mean ± SEM. *P < 0.05, **P < 0.01, ****P < 0.0001 by one-way ANOVA.*

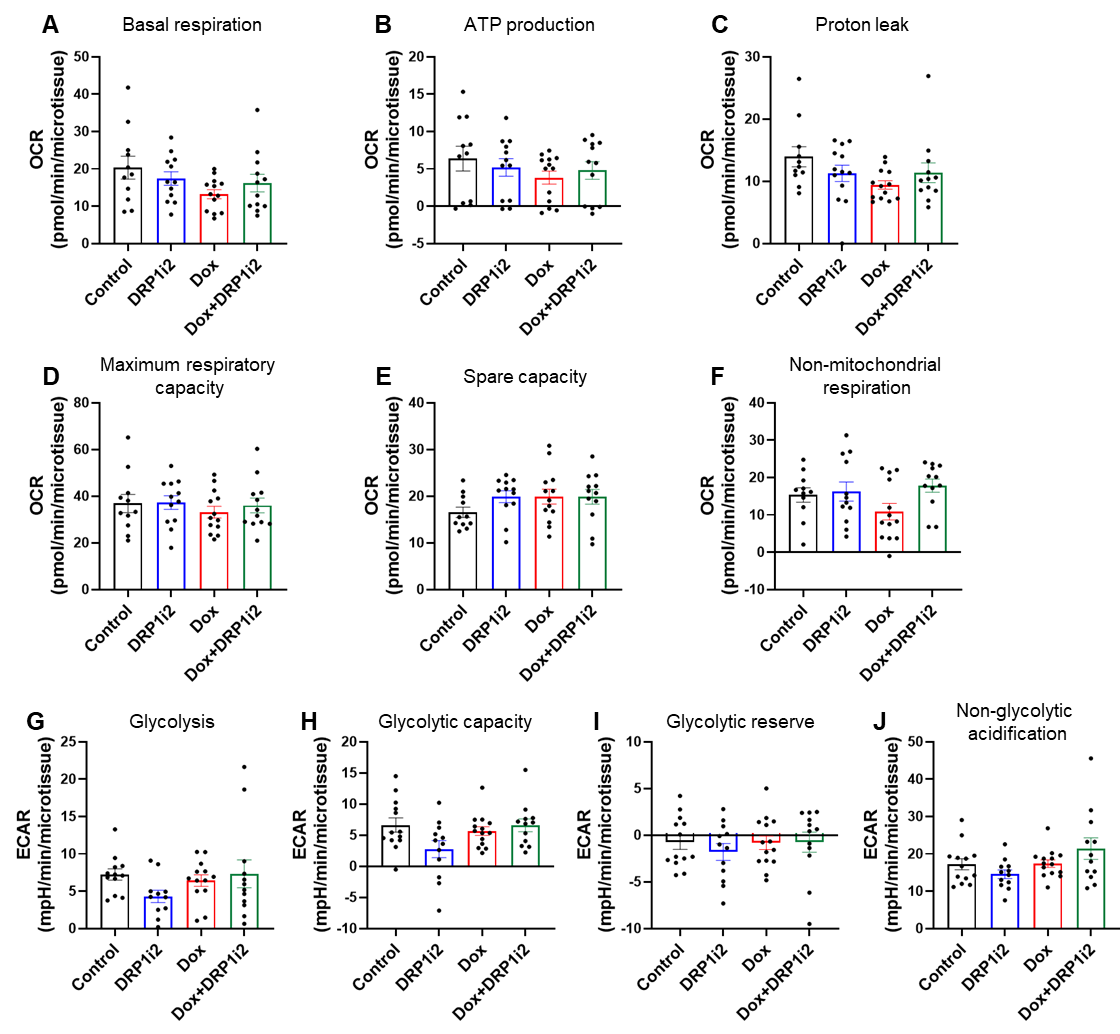

***Supplementary Figure 7. Seahorse extracellular flux analysis of energy metabolism in 3D MG63 spheroids. (A-F)*** *Mitochondrial respiration assessed by oxygen consumption rate (OCR, n = 11-13 microtissues/spheroids): basal respiration* ***(A)****, ATP‑linked respiration* ***(B)****, proton leak* ***(C)****, uncoupled respiration* ***(D)****, spare respiratory capacity* ***(E)****, and non‑mitochondrial respiration* ***(F)****.* ***(G-J)*** *Glycolytic function assessed by extracellular acidification rate (ECAR, n = 11-13 microtissues/spheroids): glycolysis* ***(G)****, glycolytic capacity* ***(H)****, glycolytic reserve* ***(I)****, and non‑glycolytic acidification* ***(J)****.*
